# ArreSTick Motif is Responsible for GPCR-β-Arrestin Binding Stability and Extends Phosphorylation-Dependent β-arrestin Interactions to Non-Receptor Proteins

**DOI:** 10.1101/2023.08.04.551955

**Authors:** András Dávid Tóth, Eszter Soltész-Katona, Katalin Kis, Viktor Guti, Sharon Gilzer, Susanne Prokop, Roxána Boros, Ádám Misák, András Balla, Péter Várnai, Lilla Turiák, András Ács, László Drahos, Asuka Inoue, László Hunyady, Gábor Turu

**Author notes:** These authors contributed equally to this work (András Dávid Tóth, Eszter Soltész-Katona, László Hunyady, Gábor Turu).

## Abstract

The binding and function of β-arrestins are regulated by specific phosphorylation motifs present in G protein-coupled receptors (GPCRs). However, the exact arrangement of phosphorylated amino acids responsible for establishing a stable interaction remains unclear. To investigate this pattern, we employed a 1D sequence convolution model trained on a dataset of GPCRs that have established β-arrestin binding properties. This approach allowed us to identify the amino acid pattern required for GPCRs to form stable interactions with β-arrestins. This motif was named “arreSTick.” Our data show that the model predicts the strength of the coupling between GPCRs and β-arrestins with high accuracy, as well as the specific location of the interaction within the receptor sequence. Furthermore, we show that the arreSTick pattern is not limited to GPCRs, and is also present in numerous non-receptor proteins. Using a proximity biotinylation assay and mass spectrometry analysis, we demonstrate that the arreSTick motif controls the interaction between numerous non-receptor proteins and β-arrestins. For example, the HIV-1 Tat Specific Factor 1 (HTSF1 or HTATSF1), a nuclear transcription factor, contains the arreSTick pattern, and our data show that its subcellular localization is influenced by its coupling to β-arrestin2. Our findings unveil a broader regulatory role for β-arrestins in phosphorylation-dependent interactions, extending beyond GPCRs to encompass non-receptor proteins as well.

## Introduction

G protein-coupled receptors (GPCRs) form one of the largest families of proteins in the human proteome, and they are prominent targets in human therapy (Hauser *et al*., 2017). Stimulation of the receptors leads to the activation of the canonical signal transduction pathways, followed by receptor phosphorylation and coupling to β-arrestin proteins (Peterson and Luttrell, 2017; Gurevich and Gurevich, 2019). β-arrestin binding results in the desensitization and internalization of GPCRs, and act as important scaffold proteins, as well. It initiates a broad range of signaling events, such as activation of mitogen-activated protein kinase (MAPK) signaling pathways, including ERK1/2, p38, and c-Jun N-terminal kinase-3, as well as that of c-Src family kinases, Akt kinase, PI3 kinase, and RhoA (Peterson and Luttrell, 2017; Gurevich and Gurevich, 2019).

Based on the duration and stability of their binding to β-arrestins, receptors can be classified into class A and class B groups. Class A receptors, including V1A vasopressin receptor, β2-adrenergic receptor (β2AR) and CB1 cannabinoid receptor, bind β-arrestin at the plasma membrane and release it quickly after internalization (Oakley *et al*., 2000; Terrillon, Barberis and Bouvier, 2004; Gyombolai *et al*., 2013). In contrast, class B receptors, such as AT1 angiotensin receptor (AT1R), V2 vasopressin receptor (V2R) and oxytocin receptor, have a stronger and more stable association with β-arrestins, which interaction can also be found at intracellular vesicles after internalization (Oakley *et al*., 2000; Oakley, Laporte, Holt and Barak, 2001). This strong and long-lasting interaction leads to a more pronounced activation of β-arrestin-dependent signaling pathways (Tohgo *et al*., 2003; Wei *et al*., 2004). GPCR–β-arrestin interaction stability depends on the phosphorylation pattern of the receptor’s C-terminus (Oakley, Laporte, Holt, Barak, *et al*., 2001). Distinct receptors have varying motifs, and the exact pattern required for tight β-arrestin binding has not been determined. Zhou et al. identified long and short phosphorylation codes (Zhou *et al*., 2017), whose presence and number in a few GPCRs, correlate with their ability to bind β-arrestins with high affinity. A common pattern in the short and long code is the presence of a PxxP motif, where P denotes a phosphorylated serine or threonine, and x any other amino acid residue, and the code is extended at the start either with a shorter Px or longer Pxx sequence, respectively. Mayer et al. analyzed the importance of the rhodopsin C-terminus serines and threonines in the tight binding of the visual arrestin. They found a pattern containing the PxxP motif similar to the short and long codes (Mayer *et al*., 2019). In recent studies, a “PxPP” motif was identified as a sequence present in many GPCR C-termini. Although this motif is important in inducing the active conformation of the β-arrestins, it does not seem to discriminate well between class A and B receptors (Isaikina *et al*., 2023; Maharana *et al*., 2023).

Although the above discussed motifs correlate with the GPCRs’ ability to interact stably with β-arrestins, they do not fully describe the sequence requirements for class A or B type interaction, and are not well defined enough to identify the exact location of the interaction. This leaves a gap in our understanding of how the β-arrestin binding to the receptors is determined at the sequence level. Moreover, although the phosphorylation-dependent interaction between GPCRs and β-arrestins is well established, for other proteins, only their phosphorylation-independent engagement with β-arrestins are usually considered. This leaves the positively-charged region of the N-domain, which is responsible for receptor C-terminus binding, as a site reserved for interactions with GPCRs only. Accordingly, here we aimed to better define the sequence requirements of the phosphorylation-dependent coupling of the GPCRs and β-arrestins, and check whether the same interaction surface of the β-arrestins could be utilized also by other non-GPCR proteins. Therefore, we established a convolutional neural network to identify the serine/threonine motif responsible for the stable β-arrestin binding using a dataset of 114 class A and class B receptors. Using only the sequence and the class information of the receptors, we identified the location of the β-arrestin coupling and could predict the interaction with high accuracy. We also found that the motif, which we term arreSTick, is present in many non-receptor proteins, which can bind to β-arrestin2 through its phosphate-binding residues. As a proof of concept, we have studied the effect of β-arrestin binding on the intracellular localization of the HTSF1 transcription factor, one of the intracellular proteins that contains the arreSTick motif. Our data show that interaction of HTSF1 with β-arrestins via the arreSTick motif can regulate its intracellular localization. The model is available on GitHub (https://github.com/turugabor/arreSTick), or it can be used online for protein prediction at www.arreSTick.org.

## Materials and Methods

### Convolutional neural network and protein predictions

The convolutional neural network model was implemented in Python 3 using the Tensorflow[2.6.1] library. The network structure is shown in Supplementary Figure 1, and the code is available at https://github.com/turugabor/arreSTick. During the training, we used either the sequence of the C-terminus or the ICL3 loop of GPCRs as an input. We set a convergence threshold of 0.8 of the area under the receiver operating characteristic curve (ROC AUC) value on the training data. The training was repeated in each round until the threshold was reached. During cross-validation, the receptor dataset was randomly divided into a training group and cross-validation set (104 vs. 10 receptors). The model was trained on the training set, and the cross-validation set was predicted based on either their training sequence or full sequence. The cross-validation was repeated 50 times, and average AUC ROC values were plotted. To visualize amino acid frequencies within the arrestin-binding regions, we trained a single network with the “grouped model” and extracted 15 amino acids from the position of the global max value in the convoluted sequence of receptors labeled as class B in the training sequence. The amino acid frequencies were calculated for each position, and the Logomaker Python library (Tareen and Kinney, 2020) was used for display. For ROC AUC curve generation, we ran a cross-validation 50 times, and each time a random set of 10 receptors were predicted. The predictions were averaged, and ROC AUC values were calculated for the predicted training set, cross-validation set, and cross-validation set using full sequences. ROC curves were calculated with different cross-validation strategies in the case of the grouped model predictions or the phosphorylation code (Zhou et al., 2017) predictions. In the case of the grouped model, cross-validations were performed similarly as described above. In each round, the hold-out receptors were predicted using the training sequences, the full sequences, or the full masked sequences. The predicted class B probabilities were averaged for each receptor, and these values were used for the ROC curve plotting. In the case of the phosphorylation codes, the total number of the short and long codes in each receptor and receptor class were used to create the plot. For the short and long codes definition we used the following regex patterns, respectively: “[S|T].[S|T][^P][^P][S|T|E|D]” and “[S|T]..[S|T][^P][^P][S|T|E|D]”. To predict all human GPCRs, we have collected receptor sequences, information, and topological data from the GPCRdb.org (Pándy-Szekeres *et al*., 2018). Circular representation of the receptors was done with the pyCirclize Python library (Shimoyama, 2022). Protein location data were collected from the Human Protein Atlas (proteinatlas.org) (Thul *et al*., 2017). Protein structure data for human proteome were collected from the AlphaFold2 website: https://alphafold.ebi.ac.uk/ (Jumper *et al*., 2021), and the sequence regions with over 70% model confidence were masked out (replaced with dummy amino acids) in proteins before the prediction. (Zhou *et al*., 2017)

### Materials and plasmid constructs

Cell culture reagents were from Thermo Fisher Scientific (Waltham, MA) and Biosera (Cholet, FR). Cell culture dishes and plates were from Greiner (Kremsmunster, AT). Plasmicin was from InvivoGen (Tolouse, FR), Coelenterazine h was obtained from Regis Technologies (Morton Grove, IL). Biotin was from SERVA Electrophoresis GmbH (Heidelberg, DE). High Capacity NeutrAvidin-Agarose Resin was from Thermo Scientific (Waltham, MA), and GFP-Trap Magnetic Agarose resin was from Chromotek (Planegg-Martinsried, DE). Anti-β-arrestin2, anti-β-arrestin1, and HRP-conjugated anti-rabbit antibodies were from Cell Signaling Technology, Inc. (Beverly, MA, USA). Phorbol 12-myristate 13-acetate (PMA) was from Sigma-Aldrich (St. Louis, MO). LC-MS grade solvents and urea were purchased from Merck (Darmstadt, DE). Mass spectrometry grade trypsin was obtained from Promega (Promega Corporation, Madison, WI, USA). Reagents used for enzymatic digestion (1,4-Dithiothreitol (DTT) and Iodoacetamide (IAA)) were purchased from Roche Diagnostics (Mannheim, DE) and Fluka Chemie GmbH (Buchs, CH).

N-terminally RLuc8-tagged wild-type and K2A (K11,12A)-mutant β-arrestin2 (RLuc8–wt-βarr2 and RLuc8–K2A-βarr2), wt-βarr2–Venus, K2A-βarr2–Venus (Tóth *et al*., 2018), βarr2–Rluc8 (Turu *et al*., 2021) and wild-type Venus-tagged β-arrestin1 (Gyombolai *et al*., 2013) have been previously described. The pmNeonGreen-C1 plasmid was kindly provided by Dr. Balázs Enyedi. GRK5–YFP was kindly provided by Dr. Marc G. Caron. Untagged rat βarr1 and βarr2 were provided by Dr. Stephen S. G. Ferguson. K10A and K11A mutations were introduced to rat β-arrestin1 by precise gene fusion PCR to create K2A-βarr1–Venus. To generate TurboID-tagged wild-type and K2A-mutant rat β-arrestin2, the coding sequence of TurboID (Branon *et al*., 2018) was synthesized in gBlock gene fragment (IDT, Coralville, IA, USA), and it was cloned into wt-βarr2–Venus or into K2A-βarr2–Venus by replacing Venus using AgeI/NotI restriction enzymes.

GRK5–FLAG was generated using annealed oligo cloning by replacing the YFP-encoding DNA sequence to that of the FLAG tag. HTSF1–mNeonGreen was produced by cloning the HTATSF1 from pCMV6-Entry (Origene, Rockville, MD, USA) vector into the pEYFP-N1 vector between AfeI/SalI restriction sites, then YFP was replaced with mNeonGreen using AgeI/NotI restriction enzymes. To create HTSF1–Venus, YFP was replaced by monomeric Venus (containing the A206K mutation, all Venus-tagged constructs used in this study harbored this monomerizing mutation). Alanine mutant form of HTSF1 (HTSF1-ST/AA: S739A, T740A, S742A, S743A, S747A, and S748A) was created by gBlock gene fragment (IDT, Coralville, IA, USA) synthesis, and it was cloned into pEYFP-N1 vector between BglII/AgeI restriction sites. After that, YFP was replaced by mNeonGreen or Venus with AgeI/NotI restriction enzymes. The L10–mRFP construct, containing the plasma membrane target sequence L10 (MGCVCSSNPENNNN, the first 10 amino acids of mouse Lck followed by polyglutamine linker), was created by replacing Venus by mRFP in L10–Venus construct (Gulyás *et al*., 2017). To generate the AT1R-Cterm–Venus construct, the coding sequence of the C-terminus of rat AT1a angiotensin receptor (residues 320–359, IPPKAKSHSSLSTKMSTLSYRPSDNMSSSAKKPASCFEVE) together with Venus from a Venus-tagged full-length receptor construct (Gyombolai *et al*., 2013) was PCR-amplified, then it was in-frame fused with DPTRSRAQASNSGGG linker to the L10 sequence by replacing mRFP in L10–mRFP. A similar strategy was used for the two following receptor C-terminus constructs. For AT1R-Cterm-TSTS/A–Venus, AT1R-TSTS/A–Venus was used as a template (Tóth *et al*., 2018), the sequence of AT1R-Cterm-TSTS/A: IPPKAKSHSSLSAKMAALAYRPSDNMSSSAKKPASCFEVE. For V2R-Cterm–Venus, the C-terminus (residues 343–371, ARGRTPPSLGPQDESCTTASSSLAKDTSS) of the human Venus-tagged V2 vasopressin receptor (Szalai *et al*., 2014) was fused together with Venus to the L10 sequence.

### Cell culture and transfection

HEK 293T cells were from American Type Culture Collection (ATCC CRL-3216 Manassas, VA). HEK 293A parent and βarr1/2 KO cells were described earlier (O’Hayre *et al*., 2017). The cells were cultured in DMEM supplemented with 10% fetal bovine serum and 1% penicillin/streptomycin in 5% CO_2_ at 37 °C. Cells were treated with plasmocin (25 μg/ml) for two weeks before the experiments. For BRET measurements, cells were transfected in suspension using Lipofectamine 2000 according to the manufacturer’s protocol and plated on white poly-L-lysine-coated 96-well plates. For co-precipitation and confocal microscopy experiments, the calcium phosphate precipitation method was used for cell transfection as described previously (Qureshi, Ahmad and Zafarullah, 2008) (Tóth *et al*., 2021). The cells were plated on poly-L-lysine-coated 10 cm plates or on µ-Slide 8 Well Ibidi (Grafelfing, DE) plates, and the medium was replaced with fresh DMEM after 6-7 hours. Cells were regularly tested for mycoplasma contamination.

### Bioluminescence resonance energy transfer (BRET) measurements

Transiently transfected HEK 293T cells were plated on poly-L-lysine-coated 96-well white-walled tissue culture plates, and the measurements on adherent cells were performed 24–28 hours after transfection. Luminescence intensities were measured using a Thermo Scientific Varioskan Flash multimode plate reader at 37 °C as described previously (Tóth *et al*., 2018). Briefly, before the measurements, we replaced the medium with a modified Kreb’s-Ringer medium (120 mM NaCl, 10 mM glucose, 10 mM Na-HEPES, 4.7 mM KCl, 0.7 mM MgSO4, 1.2 mM CaCl_2_, pH 7.4). We determined the expression of the Venus-tagged proteins by recording fluorescence intensity at 535 nm with excitation at 510 nm. After the addition of the luciferase substrate coelenterazine *h* (5 μM), we measured luminescence intensities using 530 nm and 480 nm filters. In the BRET titration experiments, luminescence intensity was measured without a filter as well in order to assess the expression of the donor-labeled construct. The BRET ratio was determined by dividing the luminescence intensities at 530 nm and 480 nm with each other (I_530nm_/I_480nm_). In the titration BRET experiments, BRET ratios were normalized to those wells in which no Venus-tagged construct was expressed. To assess the interaction between plasma membrane-targeted receptor C-termini and βarr2, cells were transfected with Rluc8-tagged wild-type or K2A-mutant βarr2 (0.001 μg/well) and with Venus-tagged receptor C termini (0.05 μg/well). After measuring the baseline BRET ratio, cells were stimulated with 100 nM angiotensin II (Ang II), 100 nM phorbol 12-myristate 13-acetate (PMA) or 100 nM arginine vasopressin (AVP), and the change of the BRET ratio (stimulated - vehicle-treated) was continuously determined. Kinetic measurements were performed in triplicate.

For the titration BRET experiments, we transfected the cells with Rluc8-tagged wild-type or K2A-mutant βarr2 constructs (0.02 μg/well) and Venus-tagged HTSF1 plasmids in increasing concentrations (0–0.2 μg/well), we also added pcDNA3.1 to keep the total amount of transfected DNA constant (0.25 μg/well). Data from all wells are shown in the BRET titration experiments.

### Confocal microscopy

β-arrestin1/2-knockout HEK 293A cells were cotransfected with L10–mRFP, HTSF1–mNeonGreen and either untagged wt-βarr2, wt-βarr1, K2A-βarr2 or pcDNA3.1 in suspension using calcium phosphate precipitation and plated immediately on poly-L-lysine-coated µ-Slide 8 well Ibidi plates. The medium was changed the next day, and 24 hours after the transfection, the cells were fixed with paraformaldehyde (4%, 15 minutes), and the nuclei were stained with DAPI. We used a Zeiss LSM 710 confocal laser scanning microscope for obtaining representative images and ImageXpress confocal microscopy for the quantification of protein localization. For the latter, 49 images per well were obtained in three channels (L10–mRFP, HTSF1–mNeonGreen, and DAPI). L10-mRFP images were used for cell segmentation, the DAPI channel for nucleus segmentation, and the mNeonGreen channel was used for the HTSF1 fluorescence determination. Images were segmented using the Cellpose Python library (https://github.com/MouseLand/cellpose), and total mNeonGreen fluorescence in the cytoplasm (cell mask minus nuclear mask) was divided with nuclear fluorescence. The applied analysis code is available on GitHub (https://github.com/turugabor/cellAnalysis).

### Immunoprecipation and immunoblot analysis of HTSF1

HEK 293T cells were transfected in suspension with plasmids encoding Venus-tagged wild-type or K2A-mutant βarr1 or βarr2. 24 h after transfection, we placed the dishes to ice and washed them with ice-cold PBS (supplemented with 1.2 mM CaCl_2_) solution. The washing step was repeated three times. Then the cells were lysed with lysis buffer (10 mM Tris-HCl pH 7.5, 150 mM NaCl, 0.5 mM EDTA, 0.5% Triton X100) supplemented with cOmplete Protease Inhibitor mixture (Roche) and Phosphatase Inhibitor Mixture 3 (Sigma). For protein cross-linking, we added 1 mM disuccinimidyl suberate (DSS) to the lysate at 37°C for 15 minutes. After that, we quenched the reaction by adding Tris-containing buffer (50 mM Tris-HCl, 50 mM L-glycine, 1.2 mM CaCl_2_, 1 mM MgCl_2_, 50 mM NaCl, pH 7.4) to the samples at a ratio of 1:10 at 4°C for 15 minutes (Saha *et al*., 2022). Samples were centrifuged at 20,800 × *g* for 10 min and the supernatants were incubated with 15 μl GFP-Trap Magnetic Agarose resin for 1 h at 4 °C. After that, the beads were washed three times with a washing buffer (10 mM Tris-HCl pH 7.5, 150 mM NaCl, 0.5 mM EDTA). We eluted the proteins from the surface of the beads using Laemmli SDS sample buffer (2x) containing 10% mercaptoethanol at 95°C for 5 minutes. Proteins were separated with SDS-polyacrylamide gel electrophoresis and were blotted onto PVDF membranes. Membranes were treated with antibodies against HTSF1 (Proteintech 20805-I-AP), followed by the treatment with HRP-conjugated secondary antibodies.

Visualization was made with Immobilon Western chemiluminescent HRP Substrate (Millipore), and fluorescence was detected with Azure c600 Western Blot Imaging System (Biosystems). The results were quantitatively evaluated with densitometry using ImageJ software.

### Affinity purification using biotin ligase

HEK 293T cells were transfected in suspension with plasmids encoding βarr2–TurboID (biotin ligase) or K2A-βarr2–TurboID and untagged α1A-adrenergic receptor. 24h after transfection, cells were serum starved for 2–4 h, then 100 μM biotin was added for 1h, and cells were stimulated with A61603 (1 μM at 37 °C) for 1 hour to allow substantial biotinylation. Reactions were stopped by placing the dishes on ice and washing them with an ice-cold PBS solution. The washing step was repeated three times. Then the cells were lysed with 2% sodium deoxycholate (SOC) buffer (Sigma), supplemented with 0.025% RapiGest (Waters), cOmplete Protease Inhibitor mixture (Roche), and Phosphatase Inhibitor salts (1 mM sodium pyrophosphate, 2 mM sodium orthovanadate, 10 mM sodium fluoride, 50 mM β-Glycerophosphate). Lysates were collected, sonicated for 45 seconds, then centrifuged at 20,800 × *g* for 10 min. Supernatants were incubated with 100 μl of High Capacity NeutrAvidin-agarose resin for 1h at 4 °C. The beads were washed three times with ice-cold supplemented SOC and once with PBS. We eluted all proteins from the surface of the beads in Laemmli SDS sample buffer (2x) containing biotin and 10% mercaptoethanol. After centrifugation, the supernatant was transferred to a new tube. To remove SDS from the protein solution, the proteins were precipitated with 1 ml 100% ethanol (at 4°C for 24 h).

### Mass Spectrometry

Enzymatic Digestion: Precipitated, air-dried samples were digested in solution using trypsin as previously described with minor modifications (Turiák *et al*., 2019). In brief, precipitated pellets were dissolved in 30 µl 8 M urea in 50 mM ammonium bicarbonate. DTT was added at a final concentration of 5 mM and incubated at 37 °C for 30 min. For alkylation, IAA was added at a final concentration of 10 mM and incubated in the dark at room temperature for 30 min. Samples were diluted 10-fold with 50 mM ammonium bicarbonate, and enzymatic digestion was performed with 1 µl 1 µg/µl trypsin overnight at 37 °C. The reaction was quenched by the addition of 1 μL formic acid. Peptide clean-up and desalting were performed on Pierce C18 spin columns (Thermo Fisher Scientific, Waltham, MA).

Nano LC-MS/MS: Mass spectrometry measurements were performed on a Maxis II Q-TOF (Bruker Daltonics, Bremen, Germany) equipped with a CaptiveSpray nanoBooster ionsource coupled to an Ultimate 3000 nanoRSLC system (Dionex, Sunnyvale, CA, USA). Samples were dissolved in 2% AcN, 0.1% FA and injected onto an Acclaim PepMap100 C-18 trap column (5 µm, 100 µm x 20 mm, Thermo Scientific, Sunnyvale, CA, USA) for sample desalting. Peptides were separated on an ACQUITY UPLC M-Class Peptide BEH C18 column (130 Å, 1,7 µm, 75 µm x 250 mm, Waters, Milford, MA, USA) at 48 °C applying gradient elution (4% B from 0 to 11 min, followed by a 120 min gradient to 50% B). Eluent A consisted of water + 0.1% formic acid, while eluent B was acetonitrile + 0.1% formic acid. MS spectra were recorded at 3 Hz, while the CID was performed at 16 Hz for abundant precursor ions and at 4 Hz for low-abundance ones. Sodium formate was used as an internal standard, and raw data were recalibrated by the Compass Data Analysis software 4.3 (Bruker Daltonik GmbH, Bremen, Germany).

Protein identification and label-free quantitation: Proteins were identified by searching against the human Swissprot database (2019_06) using the Byonic (v3.5.0, Protein Metrics Inc, USA) software search engine. First, the combined LC-MS results were searched by Byonic (30 ppm peptide mass tolerance, 50 ppm fragment mass tolerance, 2 missed cleavages, carbamidomethylation of cysteines as fixed modification, deamidation (NQ), oxidation (M), acetyl (Protein N_Term), Glu->Pyro-Glu and Gln->Pyro-Glu as a variable modification) and proteins were identified using 1% FDR limit. This protein list was used for label-free quantitation (LFQ) using MaxQuant (Cox and Mann, 2008) (software version 1.6.7), applying its default parameters except modifications listed above and each LC-MS/MS run was aligned using the “match between runs” feature (match time window 1.5 min, alignment time window 15 min). MaxQuant analysis searched only for those proteins which were identified previously by Byonic search (this makes false identification less likely), and 1% FDR was set at the protein identification level.

### Mass spectrometry data analysis

Label-free quantification of the mass spectrometry data was carried out similarly to the MaxQuant LFQ method (Cox *et al*., 2014). This method assumes that the majority of the proteome does not change between any two conditions (i.e., two β-arrestin protein partners). Peptide intensities were normalized by minimizing the sum of all squared logarithmic fold differences between samples. In the case of a protein interactome, in contrast to a full-cell proteome, some proteins might be completely absent from the sample because of the lack of interaction. Therefore, in further analysis, we calculated the protein expression ratios using all the protein peptides instead of only the paired peptides across sample pairs as described in (Cox *et al*., 2014). The missing peptide intensity values were replaced with zeros. The peptides from six experiments were pooled together, and the median log2 fold difference in intensities was taken as a difference to decrease the effect of the outliers on the average value. Differences between individual proteins in the wild-type- and K2A-mutant β-arrestin2 samples were statistically analyzed with the Wilcoxon test, and false discovery rate correction was performed with the Benjamini-Hochberg method. Protein preference to either of the β-arrestins was determined as a positive (wild-type preference) or negative (K2A preference) log2 fold difference.

### Data analysis and plotting

Unless otherwise stated, data analysis and plotting were done in Python, using Pandas, Numpy, Matplotlib, Seaborn, and Scipy libraries, and the statistical analyses were conducted using GraphPad Prism 9 Software. The prediction model is available on GitHub (https://github.com/turugabor/arreSTick).

## Results

### Identification of the serine-threonine pattern responsible for class B β-arrestin binding

The phosphorylation of the C-terminus, or in some cases the third intracellular loop (ICL3), of GPCRs is a main determinant of β-arrestin binding, and most likely, the pattern of the phosphorylation (phosphorylation barcode) determines the stability of the receptor–β-arrestin interaction (Oakley, Laporte, Holt, Barak, *et al*., 2001). We hypothesized that the specific amino acid pattern in GPCRs required for the stable interaction can be predicted with machine learning algorithms using only sequence information. By reviewing the literature, we created a receptor–β-arrestin binding-class database consisting of 114 receptors (113 GPCRs + Transforming growth factor beta (TGFβ) receptor, Figure 1A, Supplementary Table 1), which includes their β-arrestin binding properties (class B or non-class B). To build a machine learning algorithm, we opted for convolutional neural networks since these are well suited for local pattern search in 2D (e.g., images) and 1D (e.g., sequence) data structures. Convolutional networks learn kernels (i.e. assign weights for each kernel positions), which represent characteristic patterns. The analyzed sequences need to be represented (embedded) as vectors of numbers in one or more dimensions. At each point of the sequence, the dot product of the kernel and the equal-sized parts of the following sequence is calculated (Supplementary Figure 1). These products are the values of the convolution at any particular point in the sequence. Higher values mean a better match with the pattern, and we can use the maximal value along the sequence to predict the class of the receptor and the best-matching position in the sequence (Figure 1B, Supplementary Methods and see Supplementary Figure 1 for an example convolution on a single receptor).

**Figure 1.**
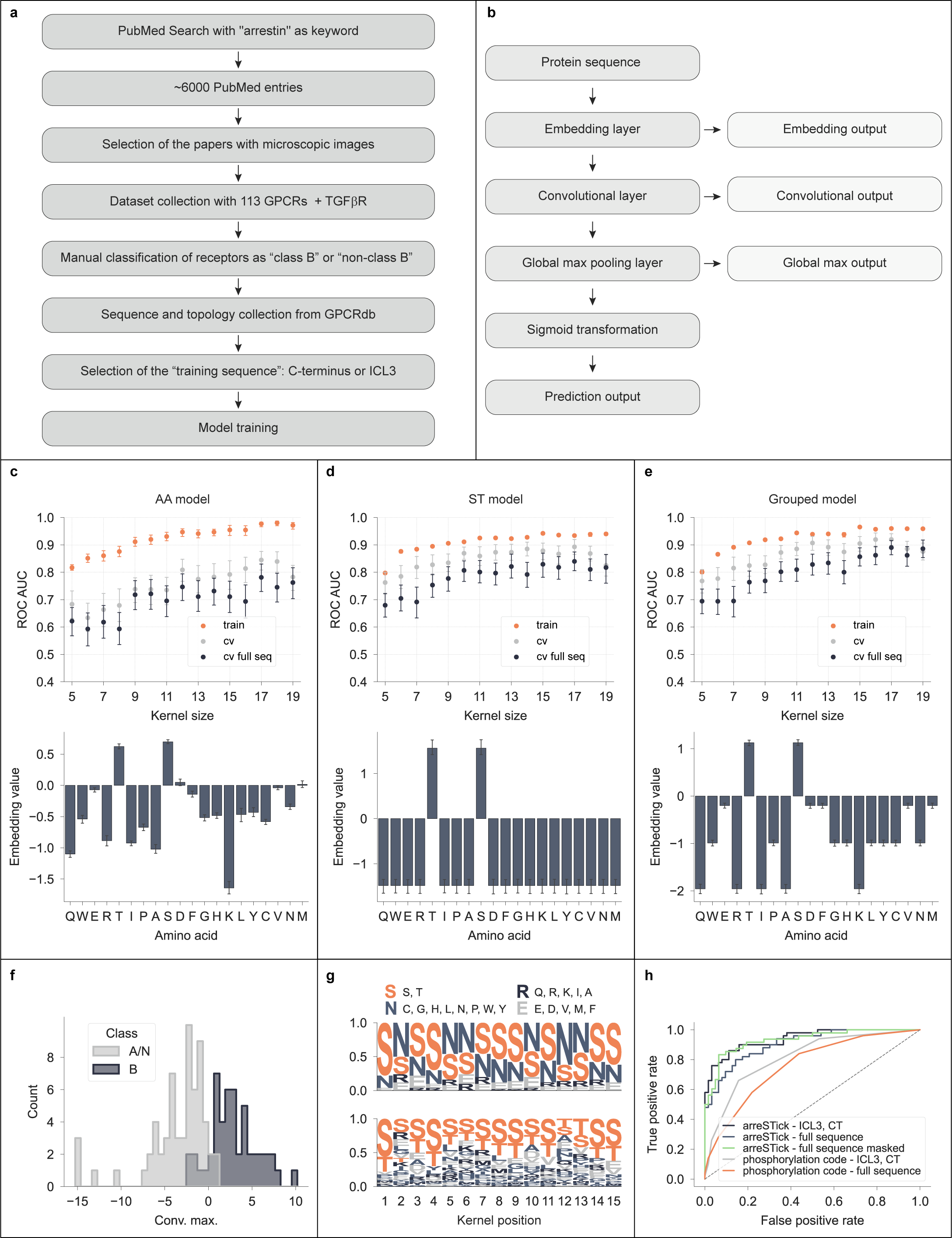
Convolutional model for prediction of the stable interaction between GPCRs and β-arrestins. A. Flowchart of the training set creation process. B. Convolutional neural network structure used in the prediction models. The protein sequence is passed to the embedding layer, where each amino acid or amino acid group is assigned a single number. The next layer is the convolutional layer which is passed to the global max pooling layer, which selects the maximum value from the convoluted values. This value is passed to a dense layer with a single neuron, from which the output through a sigmoid activation becomes a value between 0 (class A) and 1 (class B). Each hidden layer is attached to an output layer, which can be used for model interpretation. C-E. Cross-validation results (upper panels) with the convolutional models and the trained embedding values of individual amino acids (lower panels). Mean, and 95% confidence intervals are shown. Embedding values are extracted from the trained models, and the mean and 95% confidence intervals are shown on the bottom panels. F. Representative distribution of the global max values after passing the training receptor set into the model. The grouped model was trained on all receptors using the training sequences. Distribution of the global max outputs for each receptor is shown for class A and class B receptors. G. arreSTick pattern. A single grouped model was trained with all receptors in the training set, and the 15 amino-acid long sequence (arreSTick) starting at the global max convolutional value was selected for each receptor. The logo with the grouped amino acid frequencies (upper panel) and with the individual amino acid frequencies (bottom panel) are shown. The groups are labeled as follows: S (positive embedding values): [S,T]; E (neutral embedding values): [E,D,V,M,F]; R (large negative embedding values): [Q,R,K,I,A]; N: (slightly negative embedding values): all other amino acids. H. ROC curves with different cross-validation strategies using the grouped model predictions or the phosphorylation code (Zhou *et al*., 2017) numbers.

After the initial optimization steps, we opted for the simplest network structure consisting of a single convolutional layer and a single kernel, and the amino acids were represented as single floating-point numbers (Supplementary Figure 1). For most of the receptors, we used the amino acid sequence of C-termini (annotated using the gpcrdb.org API) as training data. Since some of the receptors are known to bind β-arrestins through their ICL3, for those having very short C-terminus and long ICL3 (over 80 amino acids), we used the sequence of the latter instead (Supplementary Figure 2 and Supplementary Table 2). We utilized a cross-validation (CV) strategy by dividing the entire dataset in each step into random train- and CV sets (91% and 9%), trained the model on the train set, and predicted the receptors in the CV sets. We repeated the process 50 times with random splits to gain more precise insight into the average CV accuracies. In each run, the model did not see receptors in the CV set prior to prediction. In order to find the best filter size, we also used different kernel sizes between 5 and 19. During each round, the model was trained on the C-terminal and ICL3 regions, and the cross-validation set was predicted based on their C-terminal and ICL regions or full sequences (Figure 1C-E, upper panels). After running the cross-validations, we selected an optimal kernel length and trained the model multiple times using all receptor data, and extracted the amino acid embedding values to evaluate the importance of different amino acids in the prediction (Figure 1C-E, bottom panels). Initially, we embedded all amino acids in the model (“AA model”). As shown in Figure 1C, the model performed very well on the training set, especially using kernel lengths of at least 15 amino acids. However, the CV results were substantially lower than the training performance, suggesting that the model was overfitting on the training data. Nevertheless, only serines and threonines had large positive values in the embedding, and all other amino acids had negative or nearly zero embedding values (Figure 1C, lower panel). This is in agreement with the observation that phosphorylation of certain S/T amino acids is required for strong coupling of receptors to β-arrestins (Gurevich and Gurevich, 2006). The overfitting observed in the models can likely be attributed to the high dimensionality of the data features relative to the limited number of examples used during model training. To investigate whether reducing the number of features could reduce overfitting, we grouped the amino acids and assigned the same embedding values within each group. First, we categorized the amino acids into S/T versus non-S/T amino acids (“ST model”, see trained embedding values in Figure 1D lower panel) and ran cross-validation with the model (Figure 1D upper panel). With this embedding strategy, there was less overfitting compared to the “AA model” (Figure 1D). As an intermediate embedding strategy, we also grouped the amino acids into four groups (“ST”, “EDVMF”, “QRKIA” and “CGHLNPWY” groups) based on their embedding values in the AA model since amino acids with comparable values may have similar roles in the β-arrestin binding (“grouped model”). By applying this model, the cross-validation ROC AUC values went over 0.9 and near 0.9 when the C-terminus and the full sequence was predicted, respectively (Figure 1E). For subsequent predictions, we opted for the grouped model with a kernel length of 15, as this model structure showed a good performance in the cross-validations.

The classification of receptors in the model is based on the maximal convoluted values. These max values effectively differentiate the class A receptors from class B receptors, with only a few exceptions (Figure 1F). The kernel in the trained model shows the importance of the individual positions within the sequence region, which determines whether a receptor belongs to the class B group. The sequence region within individual receptors that best matches with the kernel, when classified as class B, likely corresponds to the region that undergoes phosphorylation and binds to β-arrestin. Therefore, we named this receptor region arreSTick, referring to the sticky, phosphorylated S/T pattern in the sequence that binds to β-arrestin. 20 example kernels are shown in Supplementary Figure 3–5 from different individual trainings. For visualization of the pattern, we grouped the aminos acids into the four groups according to the grouped model, and the group frequencies in the best matching region for all GPCRs with class B binding properties using a representative kernel are shown in Figure 1G upper panel. The individual amino acid frequencies based on the same group model in these regions are shown in Figure 1G lower panel. The position of the S/T amino acids resembles some of the previously reported patterns (Zhou *et al*., 2017). Namely, one could recognize both short-(PxPxxP, e.g., positions 9–14) and long (PxxPxxP, e.g., positions 1–7) phosphorylation codes. However, the convolutional model performs better in cross-validation than the model using only the number of these reported phosphorylation codes within a sequence, particularly when the full sequence is predicted (Figure 1H). Since β-arrestins bind to the unstructured C-terminal part of the receptors, we hypothesized that excluding the structured alpha-helical parts of the receptors during the prediction could improve the predictions when full sequences are used. To implement the masking, we opted for the AlphaFold2 confidence score (Jumper *et al*., 2021; Varadi *et al*., 2022), which indicates a structured region with high confidence, when exceeding a value of 70. As anticipated, masking improves our predictions (Figure 1H). To get a closer view of the prediction of individual receptors, we convoluted the full sequences of four receptors with class B-type β-arrestin binding properties, the AT1 angiotensin receptor (AT1R), the V2 vasopressin receptor (V2R), the ACK2 receptor (ACKR2) and the complement C5a receptor 1 (C5AR1) (Figure 2.). For each receptor prediction, the models were trained without including the predicted receptor in the training set. At each point of the sequence, the convolution values (the dot product of the filter vector and the encoded sequence, starting at the given position, see Supplementary Figure 1) were calculated, and these values were sigmoid-transformed, corresponding to the probability of class B-type binding region at each point. In the cases of AT1R, V2R, and ACKR2 receptors, the predicted β-arrestin-binding regions, starting at the maximal probability values, overlap with the experimentally identified regions (Oakley, Laporte, Holt, Barak, *et al*., 2001; Wei *et al*., 2004; Dwivedi-Agnihotri *et al*., 2020; Pandey *et al*., 2021). It is important to note that the training process did not involve information about the β-arrestin binding regions of GPCRs; the models were trained using only the sequence and class information. In the case of C5AR1, the maximal convolutional value is found in a helical region, which is improbable to serve as a phosphorylation site or participate in β-arrestin binding. However, the sequence with the highest probability, outside the areas with high AlphaFold2 confidence values, coincides with the experimentally identified β-arrestin binding site (Maharana *et al*., 2023). We predicted the presence of arreSTick motifs for all GPCRs, and the identified regions are shown in Figure 3. Intriguingly, aminergic and muscarinergic receptors of the rhodopsin family of GPCRs mainly contain the arreSTick pattern in their ICL3 loop, while in most other cases, it is located in the C-terminal region. (Figure 3).

**Figure 2.**
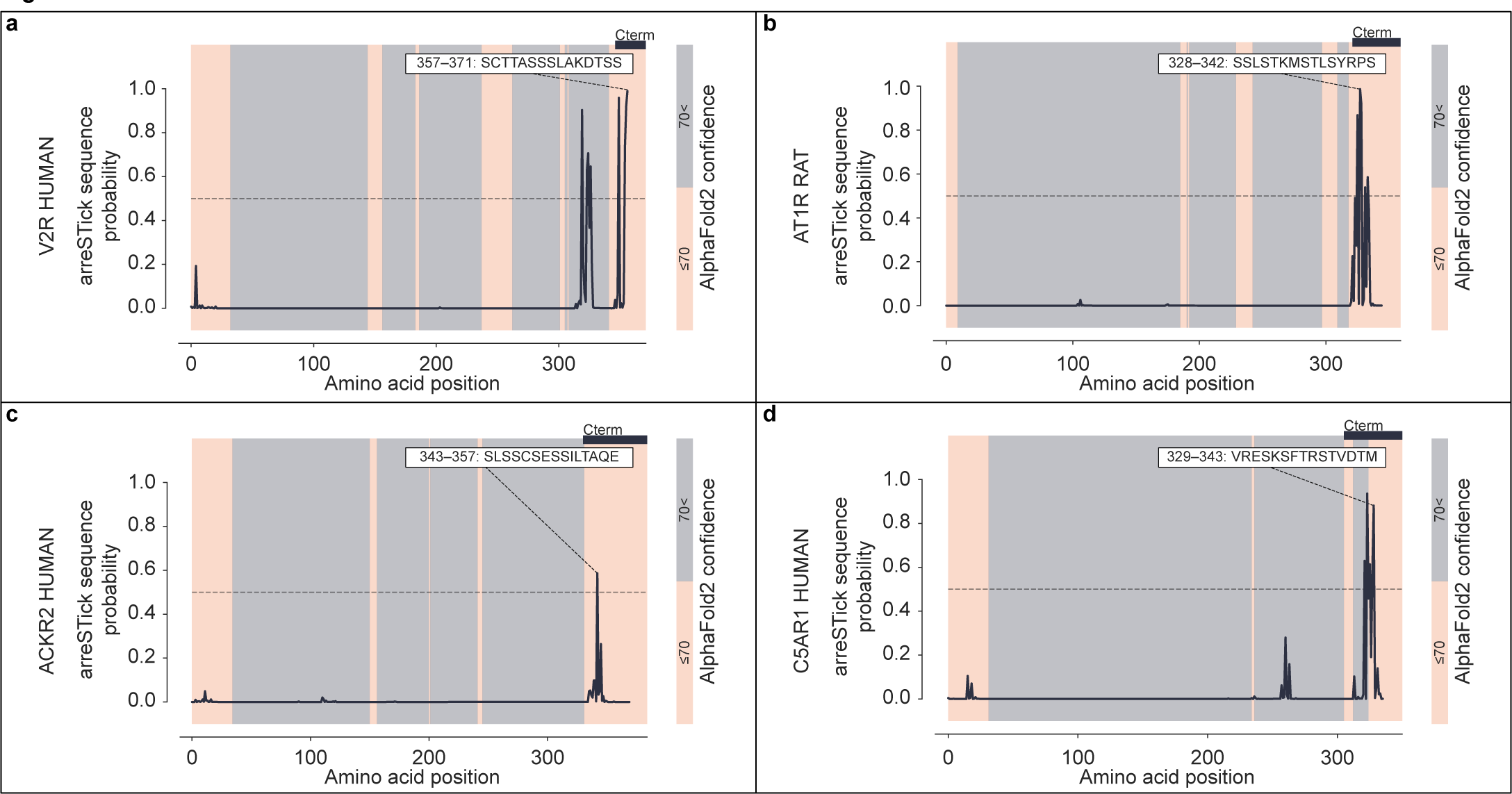
Convulution-based mapping of the arreSTick sequence for individual receptors. Full-length amino acid sequences of V2R (A), AT1R (B), ACKR2 (C), and C5AR1 (D) were passed into a trained grouped model with a kernel length of 15. Sigmoid transformation with the model’s weights was carried out on the convolutional output to get probability values at each position in the protein sequence. The arreSTick motifs, 15 amino acid-long sequence regions with the maximal likelihood of β-arrestin binding, are shown, and the AlphaFold2 model confidences (>70 or ≤70 corresponding to high and low confidence, respectively) are highlighted.

**Figure 3.**
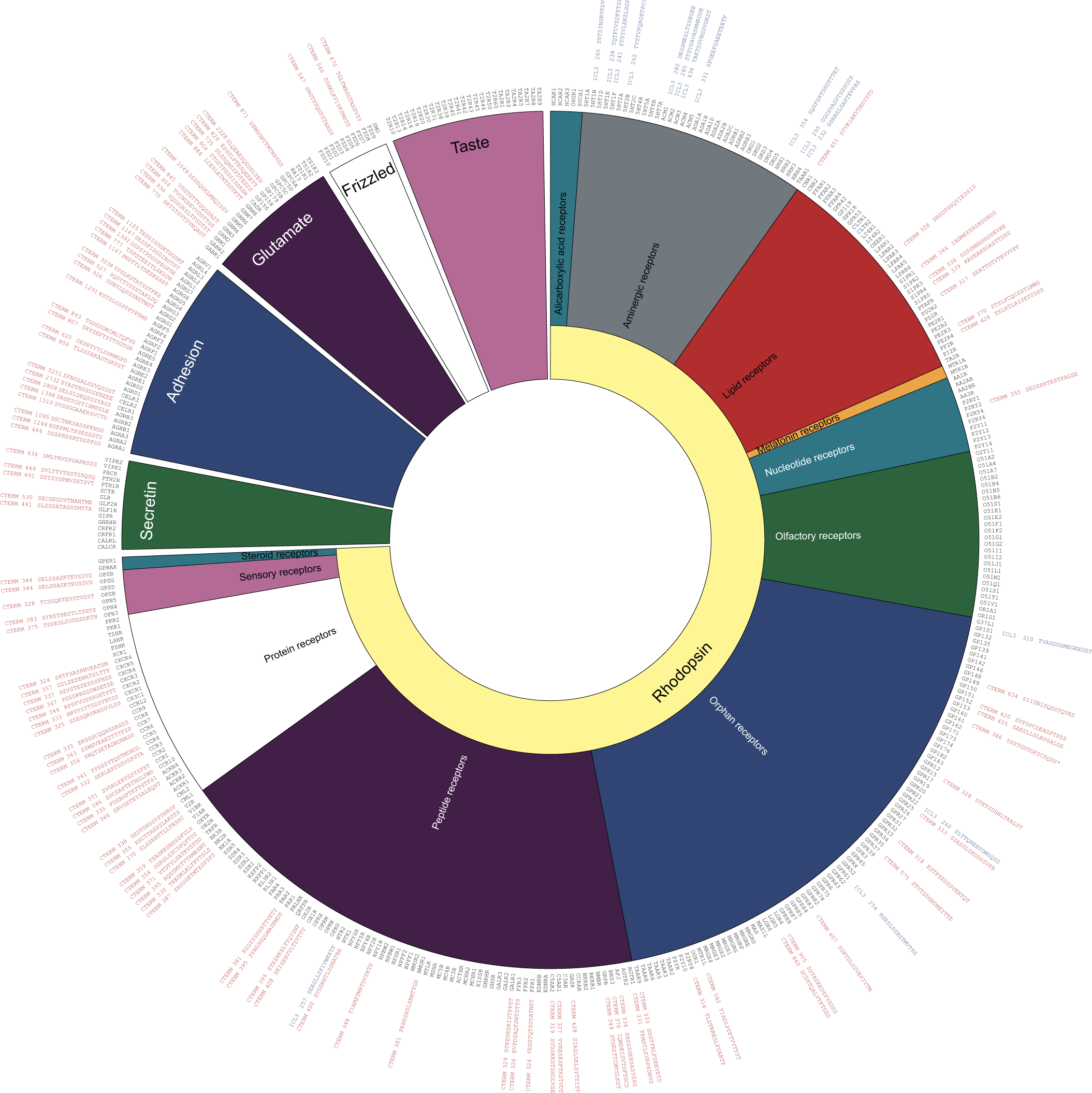
The prevalence of arreSTick in the human GPCRome. Prediction of β-arrestin binding class for all human GPCRs. The sequence information of C-termini and ICL3 regions of GPCRs was collected from gpcrdb.org. For receptors that were classified as class B, the sequence region with the highest convolutional score is presented. Receptors predicted to have arreSTick pattern in ICL3 regions are highlighted in orange. Asterisk (*) indicates that both ICL3 and C-terminus were predicted to contain the arreSTick pattern.

Phosphorylated arreSTick is sufficient for β-arrestin binding We have previously demonstrated that AT1R does not require the active state of the receptor to bind β-arrestins; phosphorylation of the C-terminus by protein kinase C (PKC) is sufficient to trigger this interaction (Tóth *et al*., 2018). Moreover, *in vitro* studies have shown that the phosphorylated C-termini of different GPCRs can also bind to β-arrestins without the involvement of the receptor core (Shukla *et al*., 2013; Kumari *et al*., 2017; Isaikina *et al*., 2023; Maharana *et al*., 2023). Therefore, we sought to determine whether phosphorylated peptides alone could interact with βarr2 in living cells. To experimentally investigate this, we designed a bioluminescence resonance energy transfer (BRET)-based setup, in which the interaction between RLuc8-labeled βarr2 and Venus-tagged C-termini of GPCRs, without the seven transmembrane structures, was assessed. To facilitate the phosphorylation of the receptor termini, they were targeted to the plasma membrane, where GPCR-phosphorylating kinases are more abundant (Figure 4A-C). The phosphorylation of the C-terminal peptide of AT1R (AT1R-CTerm) was induced by treatment with the PKC-activator PMA, while coexpression of GRK5 was applied for the V2R C-terminus (V2R-CTerm). Both strategies are known to promote activation-independent phosphorylation of these receptors (Li *et al*., 2015; Tóth *et al*., 2018; Drube *et al*., 2022). As shown in Figure 4A, the BRET signal between βarr2–RLuc8 and AT1R-CTerm–Venus increased after PMA stimulation. Angiotensin II (Ang II) had no effect, as the construct lacks the transmembrane regions responsible for Ang II binding. The binding between AT1R and βarr2 is stabilized by the interactions that we referred to as the “stability lock” in an earlier study (Tóth *et al*., 2018). A high-affinity binding is formed between amino acids K11 and K12 in the N-domain of βarr2 and the phosphorylated serine and threonine residues on the receptor C-terminus (Shukla *et al*., 2013; Isaikina *et al*., 2023; Maharana *et al*., 2023). In contrast, the K11,12A (K2A)-mutant βarr2 is incapable to establish this high-affinity interaction with GPCRs (Vishnivetskiy *et al*., 2000; Gimenez *et al*., 2012; Tóth *et al*., 2018). When we repeated the previous experiment using the phosphorylation- (and arreSTick-motif)-deficient TSTS/A-mutant AT1R C-terminus (Figure 4B), or the phosphate binding-deficient K2A-mutant βarr2 (Figure 4C), PMA treatment did not affect the BRET signal. The coexpression of GRK5, known to induce activity-independent receptor phosphorylation, led to an increase of the BRET ratio between wt-βarr2–RLuc8 and V2R-CTerm–Venus, which increase was not observed when K2A-βarr2–RLuc8 was expressed (Supplementary Figure 6). As expected, vasopressin stimulation induced no interaction since the receptor core is missing from the construct. These data show that phosphorylated GPCR C-termini without the presence of other receptor regions can be sufficient for βarr2 recruitment in living cells.

**Figure 4.**
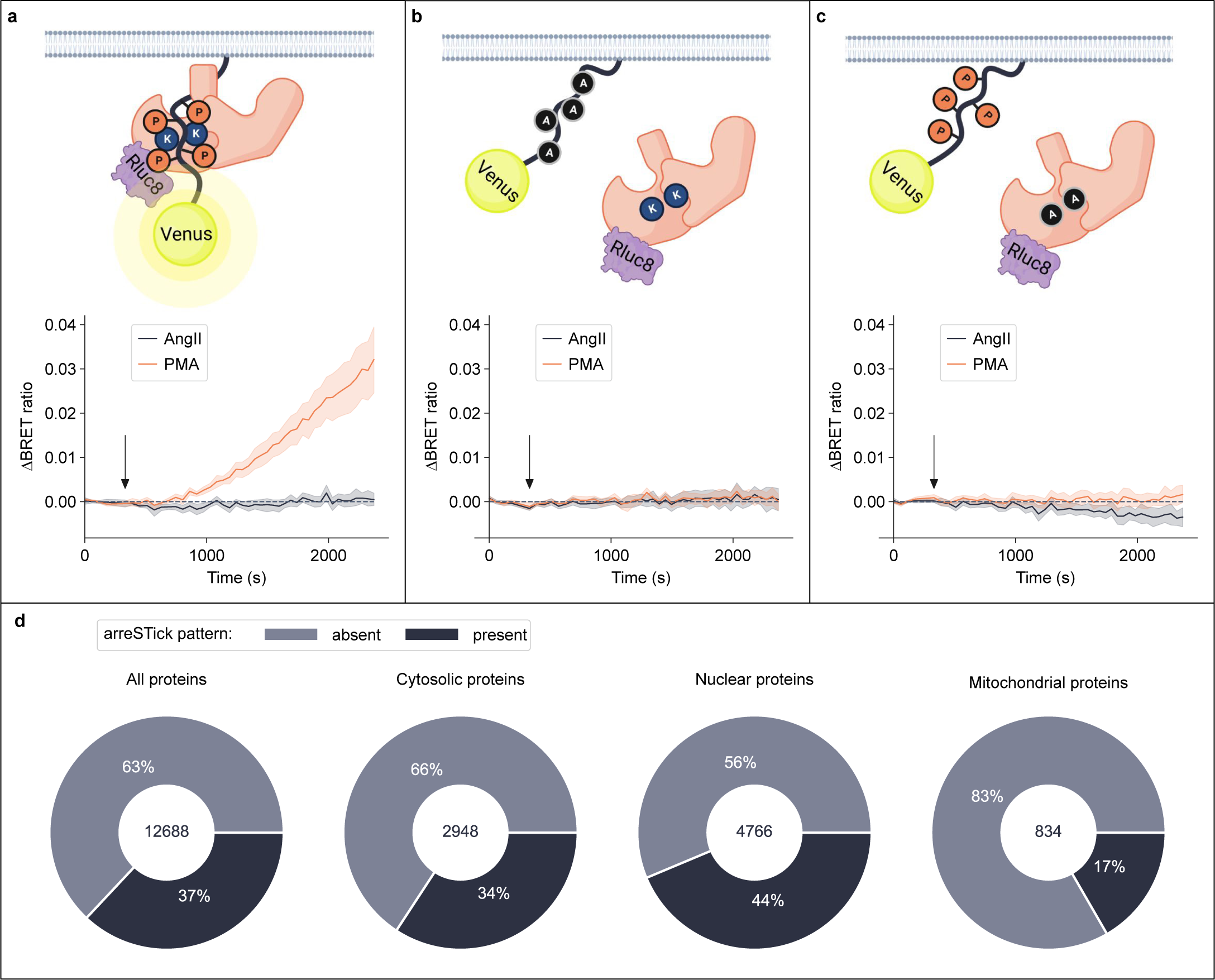
GPCR C-termini with phosphorylated arreSTick pattern can recruit β-arrestin2 even in the absence of the GPCR core region GPCR transmembrane region is not required for binding. A Kinetic BRET measurement between membrane-targeted AT1R-Cterm–Venus and wt-βarr2–RLuc8. The cells were stimulated with either 100 nM Ang II or 100 nM PMA, and the BRET signal was normalized to the vehicle-treated points. PKC stimulation by PMA led to increase of the BRET ratio, reflecting the interaction AT1R-Cterm and between βarr2. In contrast, AngII had no effect. B and C, The effect of PKC stimulation is abolished if an arreSTick motif-disrupted receptor C-terminus (AT1R-Cterm-TSTS/AA–Venus, B) or a phosphate-binding deficient βarr2 mutant (K2A-βarr2–RLuc8, were applied.The arrows show the time of treatment. Data are mean ± S.E.M., one-sample t tests were performed to statistically test whether the average changes in BRET ratio are significantly different from 0, n=3, *: p=0.0178; ns: p=0.4138; p=0.7339. D. The arreSTick pattern is present in non-GPCR proteins. Protein intracellular location data were downloaded from proteinatlas.org, and protein sequences were predicted with the grouped model. The sequence regions with predicted secondary structures (defined as AlphaFold2 confidence score above 70) were masked before prediction to decrease the number of false positives.

Cytoplasmic and nuclear proteins harboring the arreSTick pattern interact with β-arrestin2 through its cognate phosphate-binding residues Our results imply that if a protein contains an arreSTick pattern exposed to relevant kinases, the protein may also bind β-arrestins in a similar manner as GPCR C-termini. Therefore, we investigated whether the arreSTick pattern is also present in proteins other than membrane receptors. We applied the arreSTick prediction to all human proteins with known cellular locations according to the Human Protein Atlas (Thul *et al*., 2017) (Figure 4D). We excluded the protein segments that are predicted to have well-defined structures (cutoff set to AlphaFold2 confidence was 70), since these regions are unlikely to bind β-arrestins in a similar manner as the unstructured C-termini of GPCRs. Remarkably, more than 30% of all proteins were predicted to contain the β-arrestin-binding arreSTick pattern, while more than 40% of the nuclear proteins possess this potential binding site. On the other hand, mitochondrial proteins only contain arreSTick motifs in around 15%. The high number of arreStick motifs in the human proteome raises the intriguing possibility that some non-receptor proteins, if phosphorylated, may also utilize a similar phosphorylation-dependent mechanism for interacting with β-arrestins as GPCRs do.

To investigate this experimentally, we carried out a proximity labeling assay (Branon *et al*., 2018) for the interrogation of the βarr2 interactome. We designed a biotin ligase–related assay format, which exploits the fast kinetics and high activity of the TurboID ligase enzyme. We hypothesized that, if non-receptor proteins interact with β-arrestins by phosphorylated S/T amino acids, they should preferentially bind to the wild-type βarr2 over the K2A-mutant. Therefore, we used TurboID-fused wt-βarr2 or K2A-βarr2 to carry out proximity labeling in HEK 293T cells. To test the experimental setup, we first coexpressed AT1R-CTerm-Venus with wt-βarr2–TurboID or K2A-βarr2–TurboID (Figure 5A). We pulled-down the biotinylated proteins with Neutravidin beads (Tóth *et al*., 2018) and measured AT1R-CTerm–Venus fluorescence corresponding to the magnitude of the interaction between the C-terminus and βarr2. PMA treatment increased the interaction with the AT1R-CTerm–Venus only in the case of the wild-type βarr2–TurboID, confirming that this setup is able to identify proteins that bind βarr2 through a phosphorylation-dependent mechanism.

**Figure 5.**
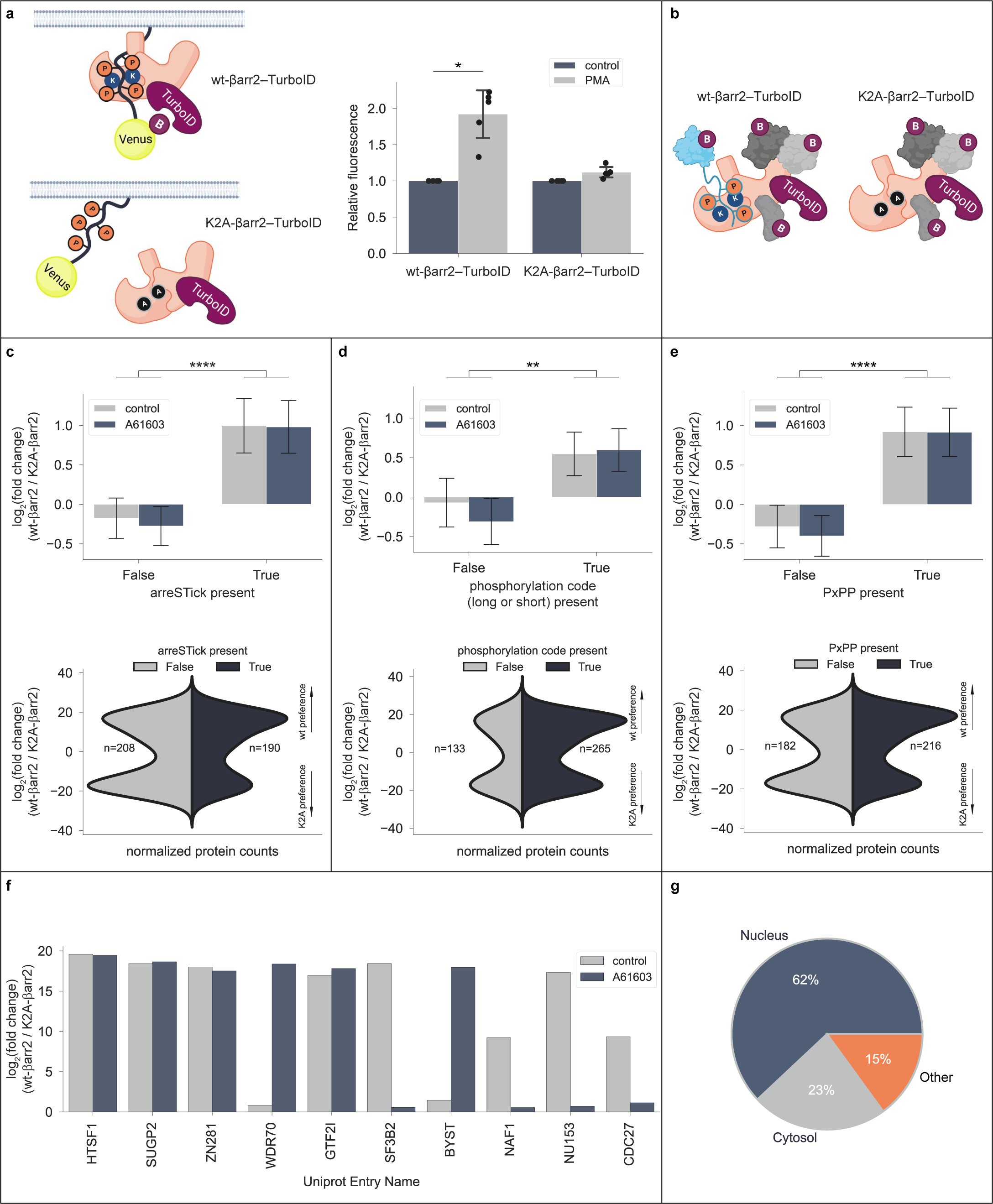
Identification of phosphorylation-dependent β-arrestin2 partners with proximity biotinylation assay. Proximity biotinylation assay-based assessment of the phosphorylation-dependent interactome of βarr2. A. Proof-of-concept measurement scheme for the βarr2–TurboID-based system using the arreSTick-containing AT1R-Cterm peptide. Wt-βarr2–TurboID or K2A-βarr2–TurboID was coexpressed in HEK 293T cells with membrane-targeted AT1R-Cterm–Venus peptide. Upon binding of TurboID-labeled βarr2, AT1R-Cterm–Venus is biotinylated, thus it can be pulled down using NeutrAvidin beads. Fluorescence of the pulled-down Venus-labeled C-termini on NeutrAvidin beads was determined. Interaction was induced by PKC stimulation using 100 nM PMA. PKC stimulation increased the amount of pulled-down AT1R-Cterm–Venus only for wt-βarr2 but not for the phosphate-binding-deficient K2A mutant. Data were normalized to the vehicle-treated conditions. Paired two-tailed t-tests were performed on the raw data, n = 5, WT-Control vs WT-PMA *, p = 0.0457; K2A-Control vs K2A-PMA n.s., p = 0.2297. B. Rationale of the proximity biotinylation assay-linked mass spectrometry experimental setup. The protein partners coupling through the arreSTick–K11/K12-βarr2 interaction (cyan-colored) are expected to be overrepresented in the interactome of the wt-βarr2, while others (proteins represented with gray color) are expected to have no preference. C, D and E. Log2 fold difference between the proteins in the interactome of the wt-βarr2–TurboID and K2A-βarr2–TurboID. HEK 293T cells were cotransfected with α1AR and wild-type or K2A-mutant βarr2–TurboID, and were stimulated with vehicle or the α1AR agonist A61603. Proteins were grouped based on the grouped model prediction (C), presence of at least one phosphorylation code (D) or the presence of the PxPP motif (E). The upper panels show the average log2 fold difference of all proteins between the wt-βarr2–TurboID and K2A-βarr2–TurboID samples, in control or stimulated samples (A61603, 100 nM). Statistical analysis was performed using two-way ANOVA, significant source of difference between the groups was the presence of the pattern (****: p < 0.0001 in c and e, **: p = 0.0092 in d), A61603 treatment induced no significant difference between the groups, and no interaction was found. The violin plots (bottom panels) show the distribution of the proteins with at least 2-fold difference between the wt-βarr2–TurboID and the K2A-βarr2–TurboID control, unstimulated samples. The width of the violins is scaled by the number of observations in each group. The number of the proteins in each group is indicated. F. Log2 fold difference of the 10 most differentially interacting proteins between the wt-βarr2–TurboID and K2A-βarr2–TurboID samples in any of the stimulated/unstimulated samples. G. Subcellular location of the proteins with arreSTick pattern and statistically significant preference for the wt-βarr2–TurboID.

Next, we applied this system to determine the entire phosphorylation-dependent interactome of βarr2s in HEK 293T cells (Figure 5B-E). ɑ1A-adrenergic receptor (ɑ1AR), a GPCR which has no detectable β-arrestin binding (Tóth *et al*., 2018), was also coexpressed in these cells to activate a broad range of cytoplasmic kinases. Therefore, half of the samples were stimulated with an α1AR-selective agonist, A61603. After isolating the biotin-labeled proteins, the samples were analyzed with label-free quantitative mass spectrometry. Altogether, we detected 1563 proteins across all samples (Supplementary Table 3). We predicted the presence or absence of the arreSTick pattern in all eluted proteins (with the exclusion of their structured regions) and investigated how the presence of the arreSTick shaped the preference of the detected proteins towards wt-βarr2 with or without α1AR and stimulation. The proteins without arreSTick regions had no preference for either of the βarr2s, however, the proteins containing this sequence had increased binding to the wt-βarr2 (Figure 5C). These results highlight the significance of the arreSTick sequence in facilitating protein binding to the positively-charged N-domain region of the βarr2. Interestingly, there was no difference between the stimulated and unstimulated samples, suggesting that the phosphorylation of arreSTick motifs for most proteins was not increased by α1AR stimulation. Next, we checked whether the proteins with previously described βarr2-binding patterns would also have preference to wt-βarr2. First, we compared those proteins that contain at least one phosphorylation code (Zhou *et al*., 2017) pattern in their sequence versus the ones without such a feature. As shown in Figure 5D, proteins with either a short or a long code showed similar tendencies as the ones with an arreSTick pattern, although the difference was less pronounced. Proteins containing the PxPP sequence, a motif previously demonstrated to play a role in the activation of β-arrestin molecules (Isaikina *et al*., 2023; Maharana *et al*., 2023), had also a preference for wt-βarr2 (Figure 5E). These data suggest that non-receptor proteins may not only bind to, but also participate in the activation ofβ-arrestins. The top 10 proteins containing the arreSTick pattern with the highest preference for wt-βarr2 are shown in Figure 5F. Next, we checked the cellular location of the wt-βarr2 preferring proteins containing the pattern according to the Human Protein Atlas (Thul *et al*., 2017). As shown in Figure 5G, most of the proteins were nuclear, with a few with cytoplasmic and other locations. These data suggest that the arreSTick motif regulates the coupling of non-receptor proteins to β-arrestins, and the coupling is dependent on the phosphoserine- and phosphothreonine-binding region of the β-arrestins, similarly to the mechanism well known in the case of GPCRs.

HIV Tat-specific factor 1 interacts with β-arrestin2 through its arreSTick region Our mass spectrometry analysis revealed that the transcription factor HIV Tat-specific factor 1 (HTSF1 or HTATSF1) exhibited the highest preference for wild-type βarr2. A previous proteomic analysis identified HTSF-1 among the proteins that immunoprecipitate with β-arr2 (Xiao *et al*., 2007), but the role of its phosphorylation in the interaction has not been studied. Our grouped prediction model predicts that HTSF1 contains an arreSTick pattern in the C-terminal part of HTSF1 (Figure 6A). This region has been previously reported to be phosphorylated (Ruse *et al*., 2008; Olsen *et al*., 2010; Hornbeck *et al*., 2012). To test the potential phosphorylation-dependent interaction of HTSF1 with both subtypes of β-arrestins, we expressed wild-type or K2A mutant Venus-labeled βarr1 or βarr2 proteins in βarr1/2 KO HEK 293A cells (O’Hayre *et al*., 2017) and performed an immunoprecipitation assay. As shown in Figure 6B, we were able to pull down endogenously expressed HTSF1 using both Venus-labeled β-arrestin1 and β-arrestin2, but not with the K2A mutants. This shows that β-arrestins interact with HTSF1, and these interactions are dependent on conserved β-arrestin residues responsible for the binding of phosphorylated serine/threonine amino acids. To further explore the specific interaction of the two proteins, we performed BRET measurements between βarr2–Rluc8 and HTSF1–Venus proteins. We opted for a titration BRET experiment to be able to distinguish the specific interaction from the non-specific energy transfer signal (Marullo and Bouvier, 2007). To verify the involvement of the predicted arreSTick pattern of HTSF1 in the binding to βarr2, we mutated the serine/threonine amino acids in this region to alanines (HTSF1-ST/AA, Figure 6C). As shown in Figure 6D, wild-type HTSF1 interacted with wild-type βarr2, reflected by the saturating BRET signal. In contrast, no BRET signal was observed when either the phosphorylation-deficient HTSF1-ST/AA-Venus or the phosphate-binding-deficient K2A-βarr2-Rluc8 mutant were expressed. These data show that the arreSTick region is involved in the interaction between HTSF1 and βarr2.

**Figure 6.**
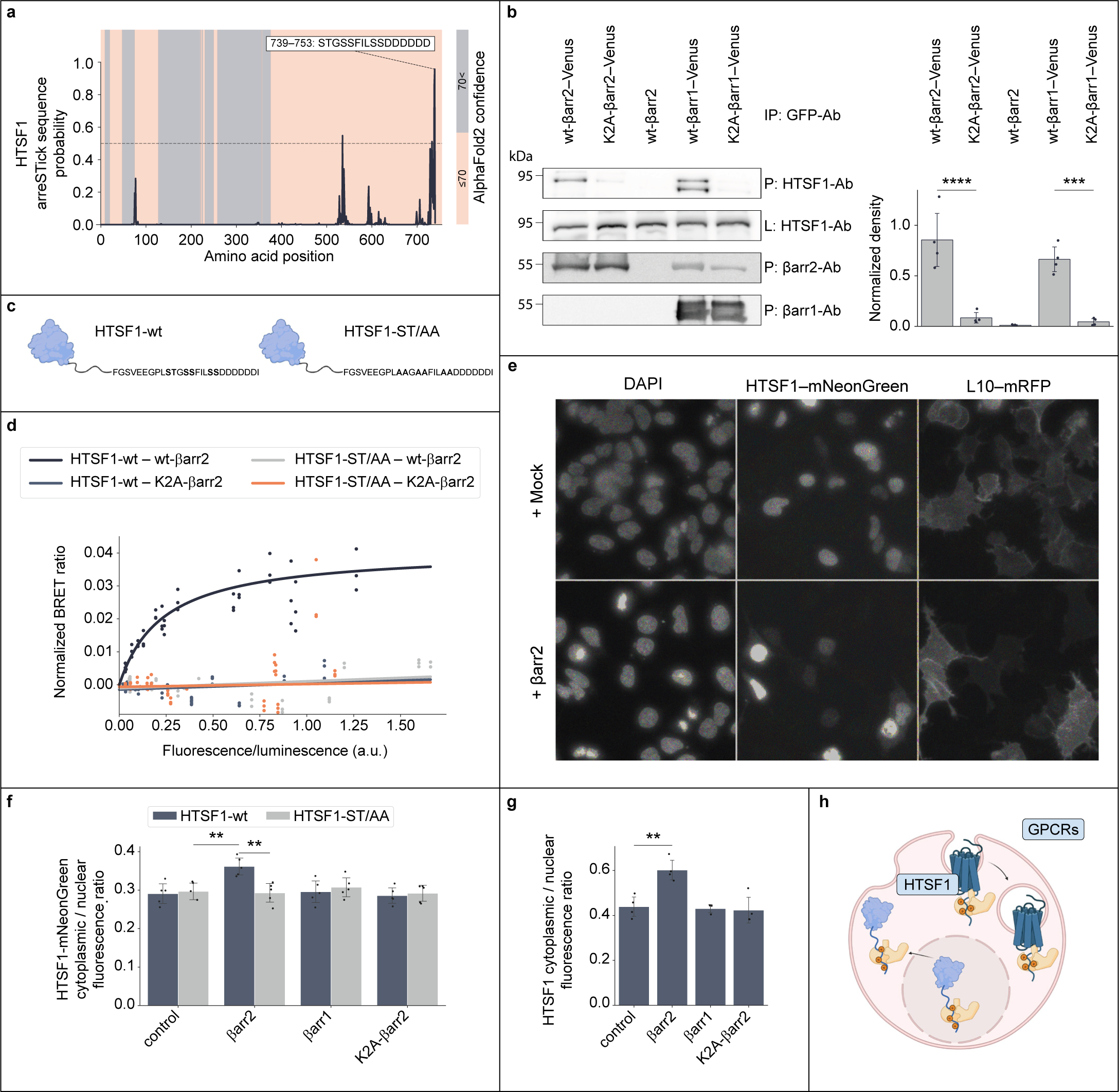
β-arrestin interacts with HTSF1 and determines its subcellular location. A. Full-length amino acid sequence of the HTSF1 was passed into the trained grouped model with a kernel length of 15. Sigmoid transformation with the model’s weights was carried out on the convolutional output to get probability values at each position in the protein sequence. The 15 amino acids long sequence region with the maximal likelihood of β-arrestin-binding shown and the AlphaFold2 model confidences (>70 or ≤70 corresponding to high and low confidence, respectively) are highlighted. B. Immunoprecipitation of the endogenous HTSF1 with overexpressed wt-βarr2–Venus, K2A-βarr2–Venus, wt-βarr1–Venus and K2A-βarr1–Venus. Non-labeled wt-βarr2 was used as a control (middle lane). The precipitation was carried out with an anti-GFP antibody, and the blots were stained with an anti-HTSF1 antibody (first row left). Lysates were stained for HTSF1 (second row), βarr2 (third row), or βarr1 (bottom row). Densities from 4 independent experiments are shown on the right, ****: p<0.0001, ***: p<0.001. C. Sequence of the arreSTick-containing C-terminal region of HTSF1 and the mutations in the HTSF1-ST/AA construct. D. BRET titration experiments with coexpressed Venus-labeled HTSF1 and RLuc8-labeled β-arrestin2. The transfected amount of βarr2–Rluc8 was held constant while the expression of Venus-labeled HTSF1 was continuously increased, and the detected BRET ratios were normalized to the samples with only βar2–Rluc8 expression. BRET ratios of individual wells from 3 independent experiments are shown; a one-site specific binding curve was fitted on the HTSF1-wt+wt-βarr2 data. Since this equation resulted in ambiguous fits for the other conditions, simple linear regression was used in these cases. E. Representative confocal images of βarr1/2 KO HEK 293A cells coexpressing wild-type HTSF1–mNeonGreen and the plasma membrane marker L10–mRFP with (bottom) or without (top) untagged βarr2. Nuclei were stained with DAPI. The images were gamma corrected with a value of 0.5 for better visualization of the low cytoplasmic HTSF1. F-G. Quantification of the distribution of HTSF1 in βarr1/2 KO HEK 293A cells. HTSF1 was either labeled with mNeonGreen (F, **: HTSF1-wt control vs. HTSF1-wt βarr2 p=0.0037; HTSF1-wt βarr2 vs. HTSF1-ST/AA βarr2 p=0.0053), or the endogenous HTSF1 was immunostained (G, **: p=0.0032), and cytoplasmic/nuclear fluorescence ratios were calculated. Ratios from 3 independent experiments are shown, the results were analyzed by two-way ANOVA (βarr2 and mutation) and Tukey’s post-hoc test was applied. H. Model of the arreSTick function in protein-protein interactions involving βarr2. In the case of GPCRs, βarr2 binding leads to receptor internalization and compartment change; the arreSTick motif stabilizes the interaction. In the case of HTSF1, the binding to βarr2 through the arreSTick motif also leads to compartment change, in this case, from the nucleus into the cytoplasm.

### β-arrestin2 regulates the intracellular location of HTSF1

βarr2 protein harbors a nucleus export signal, raising a possible function of this interaction in the regulation of the intracellular location of the HTSF1 protein, similarly to that which was already reported for some other βarr2-partner proteins, such as Mdm2 and JNK3 (McDonald *et al*., 2000; Wang *et al*., 2003). In contrast to βarr1, βarr2 harbors a nuclear export signal

(Scott *et al*., 2002), which may aid the nucleocytoplasmic translocation of partner proteins. Therefore, we hypothesized that βarr2 may play a similar role in the case of HTSF1, and the nucleocytoplasmic transport may be dependent on the interaction between the arreSTick pattern and the positive charges in βarr2 that bind to phosphorylated peptides. To investigate this, we have coexpressed HTSF1-wt–mNeonGreen or HTSF1-ST/AA–mNeonGreen with either wt-βarr2 or K2A-βarr2 in βarr1/2 KO cells. In cells expressing only the HTSF1-wt–mNeonGreen, HTSF1 localized mainly to the nucleus. When we coexpressed wt-βarr2, HTSF1 localization shifted toward the cytoplasm (Figure 6E). For the automated and unbiased quantification of the subcellular localization of HTSF1–mNeonGreen in each individual cell, we took advantage of ImageXpress high-throughput fluorescence microscopy and a deep learning-based cellular segmentation algorithm, Cellpose (Stringer *et al*., 2021) (Figure 6F and Supplementary Figure 7A). As shown in Figure 6F, HTSF1-wt–mNeonGreen cytoplasmic-to-nuclear fluorescence ratio was increased upon coexpression of wt-βarr2 with HTSF1-wt–mNeonGreen. However, this increased cytoplasmic localization was not detected with wt-βarr1, K2A-βarr2 or when HTSF1-ST/AA–mNeonGreen was overexpressed. Similarly, if we immunostained the endogenous HTSF1 (Supplementary Figure 7B), the cytoplasmic/nucleus fluorescence ratio increased when wt-β-arrestin2 was coexpressed but not when β-arrestin1 or K2A-β-arrestin2 (Figure 6G). The data indicate that the interaction between the arreSTick motif and βarr2 modulates the intracellular location of HTSF1, suggesting that non-receptor proteins can undergo phosphorylation-dependent regulation by β-arrestins similar to GPCRs.

## Discussion

Over the past two decades, it has been established that GPCRs display varying affinities for one of their main interacting partners, β-arrestins. The strength of this binding dictates the duration of β-arrestin coupling to GPCRs and substantially influences receptor trafficking and signaling (Oakley *et al*., 1999). GPCRs interact with β-arrestins via their cytoplasmic side of the transmembrane regions (also referred to as “core interaction”) and their phosphorylated C-terminus (Shukla *et al*., 2014; Kumari *et al*., 2016). It seems that the interaction between the GPCR C-terminus and the positively-charged groove on the N-domain of β-arrestins determines the binding stability and induces the active conformation of β-arrestins as well (He *et al*., 2021). However, the precise requirements for stable interaction remain elusive. Zhou et al. have identified a short and a long code in the GPCRs’ C-terminus, which, in a small set of the receptors, correlated well with the receptor–β-arrestin interaction stability. However, in our extended dataset with 114 examples, the presence and/or the number of these phosphorylation codes alone did not fully predict the β-arrestin-binding property of the receptors, especially when analyzing entire receptor sequences. Therefore we wanted to refine the sequence requirements responsible for stable binding between these two proteins by applying convolutional neural networks. Convolutional neural networks are well-suited for the classification of data with spatial information, such as images or sequences. Convolutional networks can employ multiple kernels and a deep structure, allowing them to recognize increasingly more complex structures at each layer. In our neural network models, we observed that increasing the number of kernels, hidden layers or the embedding dimension of amino acids quickly improved the training accuracy to 100%. However, this came at the cost of significantly lower cross-validation accuracies, a phenomenon known as overfitting in machine learning. This was anticipated, given the relatively small number of training examples. Therefore, we opted for a simpler network structure featuring only one hidden layer, one-dimensional embedding, and a single convolutional filter. Additionally, we incorporated a global max pooling layer, ensuring that no further pattern processing occurred beyond the convolutional layer and that information from distant regions in the sequence was not combined. Furthermore, we reduced the number of amino acid features by grouping amino acids based on their embedding values in the “AA model”. Despite its simplicity, this model achieved over 0.9 ROC AUC values in our cross-validation strategy. An additional advantage of using a single kernel and convolutional layer is the improved interpretability of our model, which allows greater insight into the classification process. For instance, with this structure, we can pinpoint the exact region based on which the model classifies the receptors and explore the kernel for hints about which positions in the region are more important. While the exact kernels may differ in the same model structure when trained multiple times due to randomly initialized kernel weights, we observed only minor variations when the training was repeated on several occasions (Supplementary Figure 3–5). It should be noted that the kernels are not composed of binary values (e.g., zeros and ones), so a simple S/T code cannot be derived from the kernel. However, some amino acid positions seem to have greater importance than others (Figure 1G), corresponding to positions that may be sterically available for binding to the positively charged amino acids on the N-domain of β-arrestins (Zhou *et al*., 2017; Maharana *et al*., 2023). Interestingly, the kernels suggest that the position of the phosphorylated amino acids required for the strong interaction is not strictly determined, allowing some variations. Since there are a number of possible positively charged amino acid partners in β-arrestins (Maharana *et al*., 2023), these variations may lead to slightly different β-arrestin binding and active conformations, consistent with the “barcode theory” (Nobles *et al*., 2011). Variations of the phosphorylation-specific micro-locks may lead to distinct β-arrestin activations and signaling outcomes (Sente *et al*., 2018).

Our model classifies the receptors based on the sequence which matches the learned kernel the most. Therefore, it is likely that this sequence encodes the region that directly binds β-arrestins. In accordance with that, the identified arreSTick motifs in GPCRs overlap very well with the experimentally determined β-arrestin-binding regions (Figure 2), despite not using any information on the coupling sites during the training. Compared to the variability in the kernels across different model trainings, there was even less difference in the embedding values of the amino acids (low confidence intervals in Figure 1C–D bottom panels vs. the variability of the kernels in Supplementary Figure 3–5). This suggests that the amino acid composition of the regions is more important than the exact location of the amino acids. The importance of the serine-threonine amino acids, which are phosphorylatable, is evident. However, the embedding values of other amino acids were also consistent (Figure 1C) across model trainings. Interestingly, certain amino acids seem to have a negative impact on the stable interaction. Although this is predictable in the case of the positively-charged arginine and lysine or the amino-group-containing glutamine, it might be surprising in the case of alanine. While the exact reason for this is unknown, it is possible that alanines interfere with the phosphorylation of nearby residues. Indeed, alanine is not among the preferred amino acids within the phosphorylation motif for GRKs (Johnson *et al*., 2023). It is also noteworthy that glutamate and aspartate, which are negatively charged and have been suggested as part of the phosphorylation code (Zhou *et al*., 2017), had a neutral impact on the classification within our binding motif (Figure 1C). Although our model achieved good cross-validation results, some receptors were misclassified. For example, the CB1 cannabinoid receptor, a known class A receptor (Gyombolai *et al*., 2013), was frequently classified as class B in more than half of the cases, while the B2 bradykinin receptor, a classical class B receptor (Simaan *et al*., 2005), was usually predicted as a class A receptor (Figure 3). These data suggest that other amino acid features in the receptor sequence might not be captured by our model or that the training set contains misclassified receptors that interfere with the training process. Indeed, some receptors only show class B-type β-arrestin-binding with some ligands but not with others (Rajagopal *et al*., 2010), suggesting that not all agonists can activate the receptor kinases in the same way. The refinement of the model to reach even higher precision would need more receptor examples to avoid overfitting even with more complex models (e.g., more detailed embeddings or more complex network structures). Nevertheless, our model shows a substantial ability to distinguish between class A and B receptors.

While the primary role of β-arrestins is believed to be the regulation of GPCRs, many GPCR-independent functions have also been described (Ma and Pei, 2007; Gurevich and Gurevich, 2014). β-arrestins are considered to bind to GPCRs in a phosphorylation-dependent manner, and they recruit and regulate other signal-transduction proteins through their other interaction sites (Peterson and Luttrell, 2017). We hypothesized that if other proteins contained the arreSTick pattern, they may also bind to β-arrestins in a similar manner as GPCRs, suggesting the potential existence of a novel β-arrestin-dependent regulatory mechanism. Therefore, we investigated whether similar arreSTick patterns could be identified in other proteins within the proteome. Unexpectedly, a substantial proportion of non-GPCR proteins possess patterns that may be capable of binding to β-arrestin with high affinity, given they are phosphorylated (Figure 4D). Indeed, the presence of the arreSTick pattern in non-GPCR proteins led to a preference for the wild-type βarr2 over the K2A-mutant in the βarr2 interactome. The K2A mutant of βarr2 lacks the ability to bind to the phosphorylated C-termini of AT1R and V2R, thus serving as a valuable tool to distinguish between βarr2 partners that are dependent on the phosphorylated arreSTick pattern for the interaction and those that are not. However, the pattern most likely needs to be phosphorylated for the interaction to occur, given the phosphorylation dependence of GPCR-β-arrestins interactions. In our experimental setup, we should have only detected those proteins which are either constitutively phosphorylated in HEK 293 cells or whose phosphorylation can be induced by α1AR stimulation. For this reason, not all partners harboring arreSTick patterns are anticipated to bind to βarr2 in this assay, but we expect enrichment of true positive binding partners in the wt-βarr2 interactome.

We further analyzed the binding of the HTSF1 protein to β-arrestin because this protein displayed the highest preference for the wild-type βarr2 over the phosphate-binding deficient K2A-mutant. HTSF1 is a protein initially recognized as protein regulating the human immunodeficiency virus type 1 (HIV-1) gene expression (Zhou and Sharp, 1996; Miller *et al*., 2009; Hulver *et al*., 2020). It might be also involved in the formation of metastases (Chang *et al*., 2021). The phosphorylation of the HTSF1 in the arreSTick region has been previously reported (Ruse *et al*., 2008; Olsen *et al*., 2010; Hornbeck *et al*., 2012). We found that the binding of β-arrestins to HTSF1 was dependent on both the arreSTick pattern and the K11 and K12 amino acids, suggesting an interaction similar to that observed with GPCRs. Interestingly, the binding of βarr2 to HTSF1 induced its translocation from the nucleus to the cytoplasm, implying a potential role in the regulation of this transcription factor. This translocation appears analogous to the role of β-arrestins with GPCRs, wherein the phosphorylated protein is removed from its primary site of action and transported to another compartment (Figure 6H). In the case of GPCRs, the active phosphorylated receptors bind to β-arrestins and undergo internalization into intracellular vesicles while being desensitized during this process. In the case of HTSF1, the nucleus is the primary site of action, and the phosphorylated HTSF1 might be transported into the cytoplasm when coupled to βarr2. Notably, βarr1 was not able to induce a similar translocation, consistent with the absence of a nuclear export signal on this protein (Wang *et al*., 2003).

In summary, we described the amino acid pattern required for stable interaction between the C-terminus of GPCRs and β-arrestins. The identified region is present not only in GPCRs but also in other proteins, in which it regulates protein-protein interactions. These findings suggest that the role of β-arrestins in regulating phosphorylated proteins may be more extensive than previously recognized.

## Acknowledgments

The technical assistance of Eszter Halász, Ilona Oláh, and Kata Szabolcsi is greatly appreciated. Figure drawings were created with BioRender.com.

---

Supplementary Table 1

Receptor dataset that was used for the model training.

Supplementary Table 2

Receptor sequences that were used for the training.

Supplementary Table 3

All proteins detected by mass spectrometry with their average log2 fold differences, p values and presence of.

**Supplementary Figure 1.**
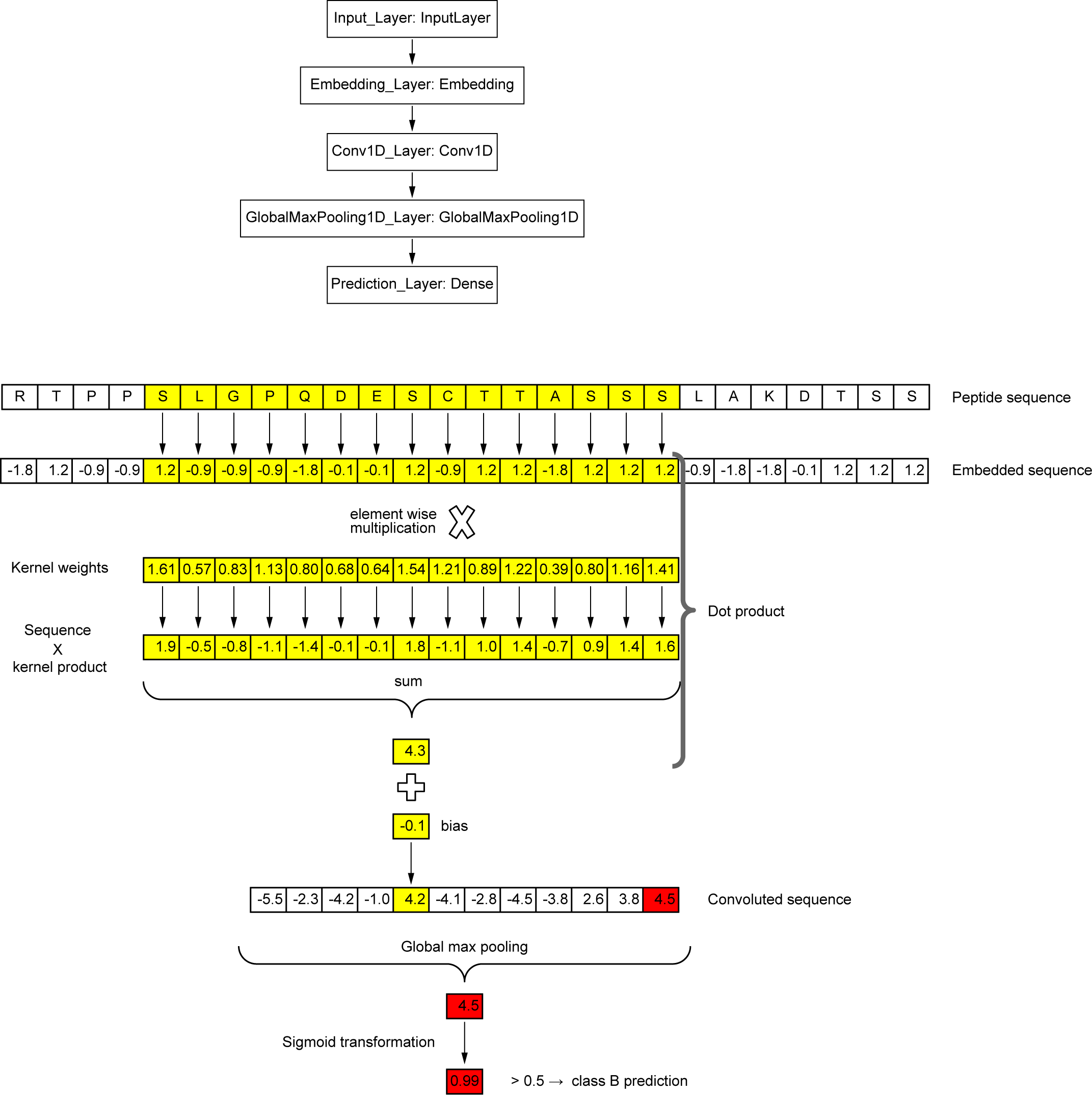
Convolutional neural network structure. The structure of the grouped model (upper) and an example convolution of the V2R. To process the input sequence, in the first step, it is passed through the embedding layer, which converts the amino acids to numerical values. Next, a 1D convolutional layer applies a kernel that slides through the sequence and calculates the dot product between the kernel and each 15-amino-acid segment. The output of this convolutional layer is then passed to a GlobalMaxPooling layer, which selects the maximum value from the previous step. Finally, the resulting value is transformed by a sigmoid function to generate the prediction.

**Supplementary Figure 2.**
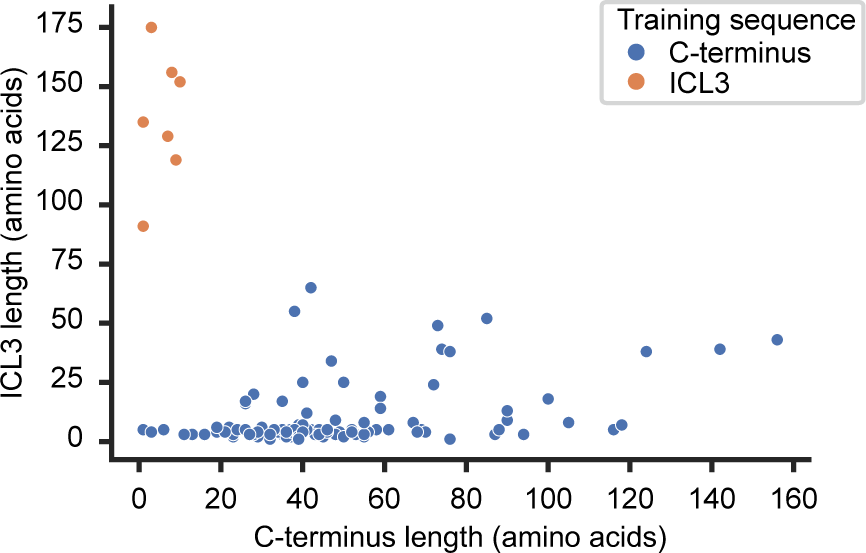
Amino acid lengths of the GPCRs’ C-termini and ICL3 regions. The sequence lengths of the C-terminus and the ICL3 region are plotted for the receptors. Each dot represents a receptor in the training set. In the case of receptors with very short C-terminus and long ICL3 region (red dots), the ICL3 sequence was used for the training instead of the C-terminus.

**Supplementary Figure 3.**
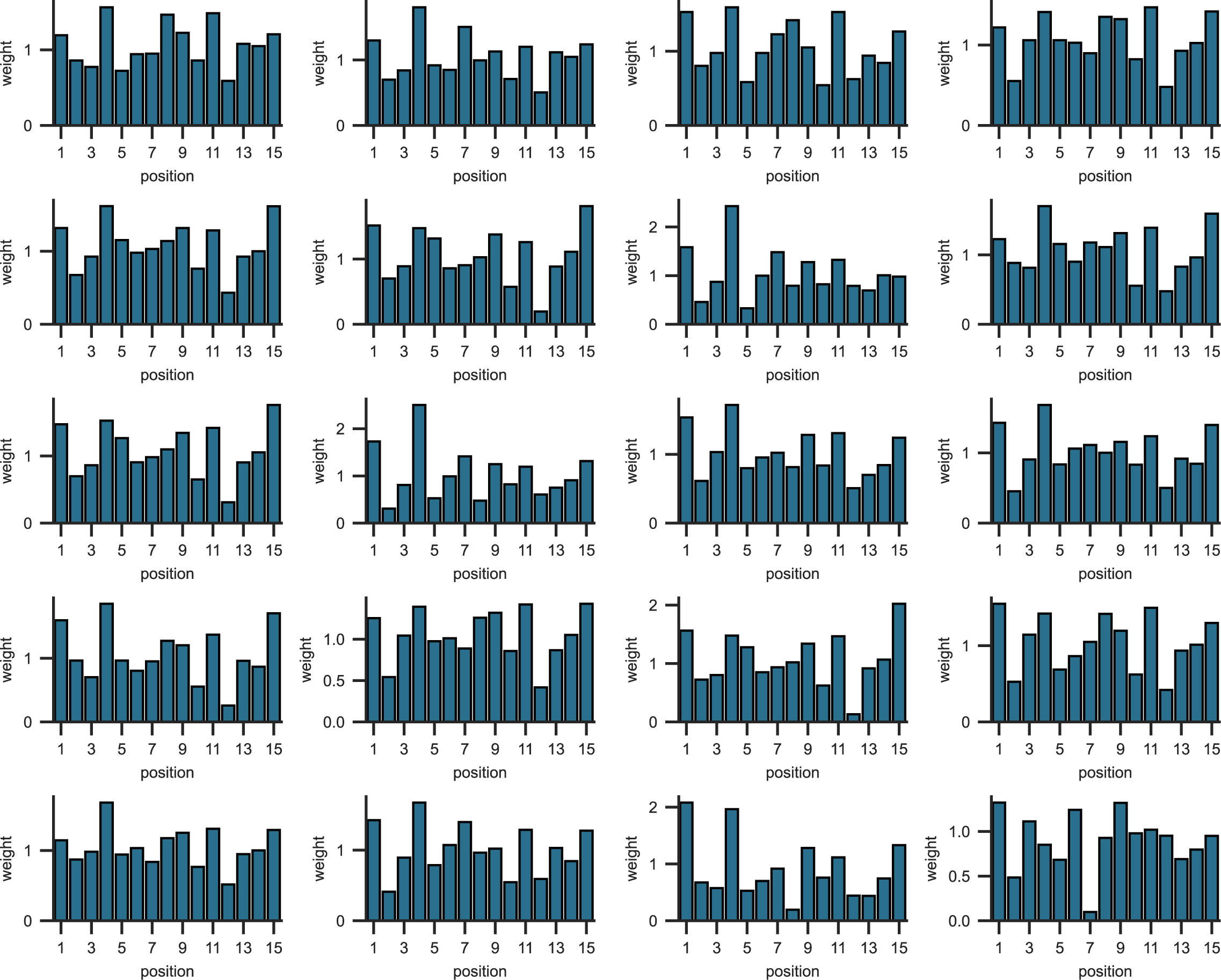
Kernel examples from “AA” model training repetitions. The “AA” has undergone 20 training runs with random initiations, and the resulting learned kernels are displayed.

**Supplementary Figure 4.**
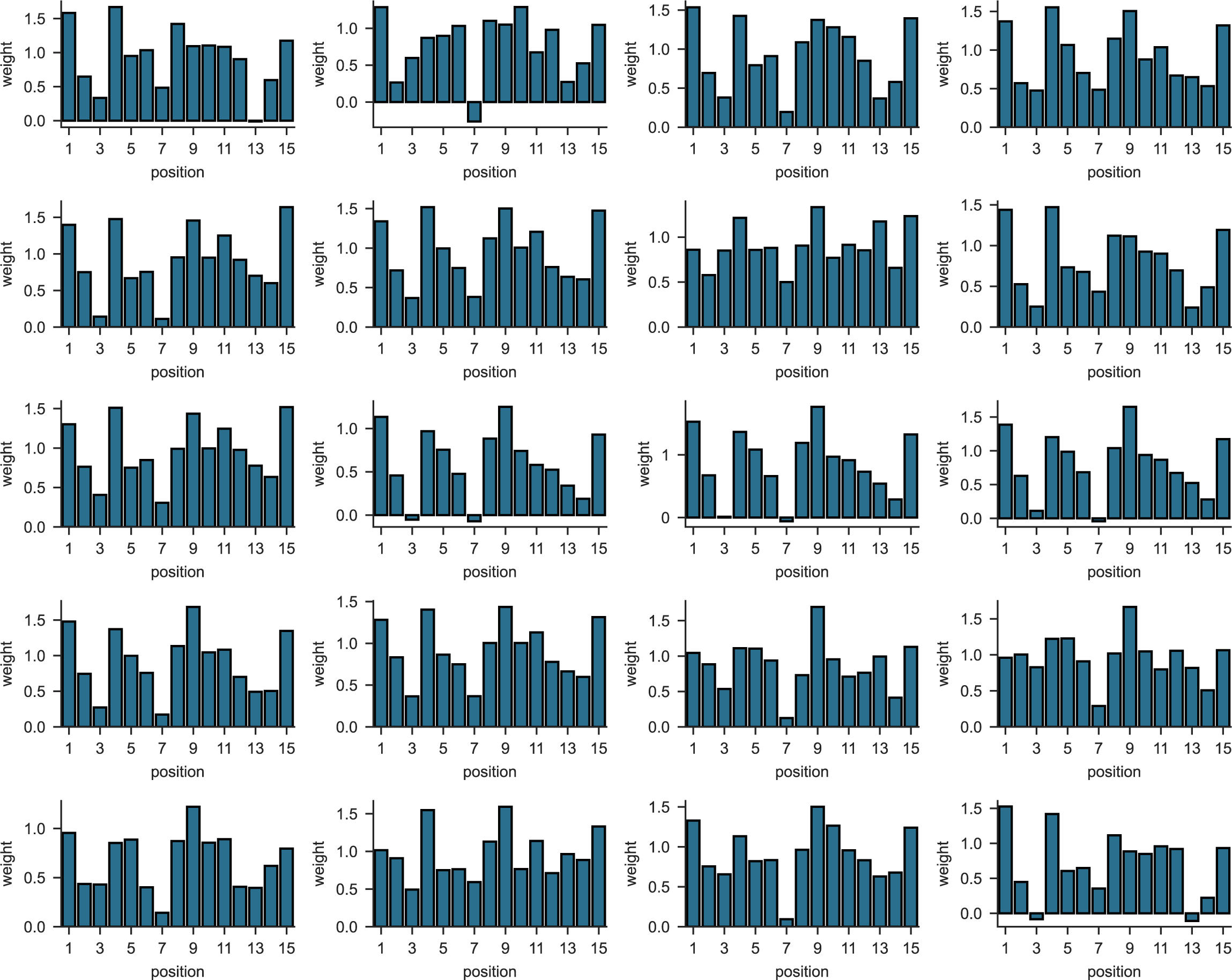
Kernel examples from “ST” model training repetitions. The “ST” has undergone 20 training runs with random initiations, and the resulting learned kernels are displayed.

**Supplementary Figure 5.**
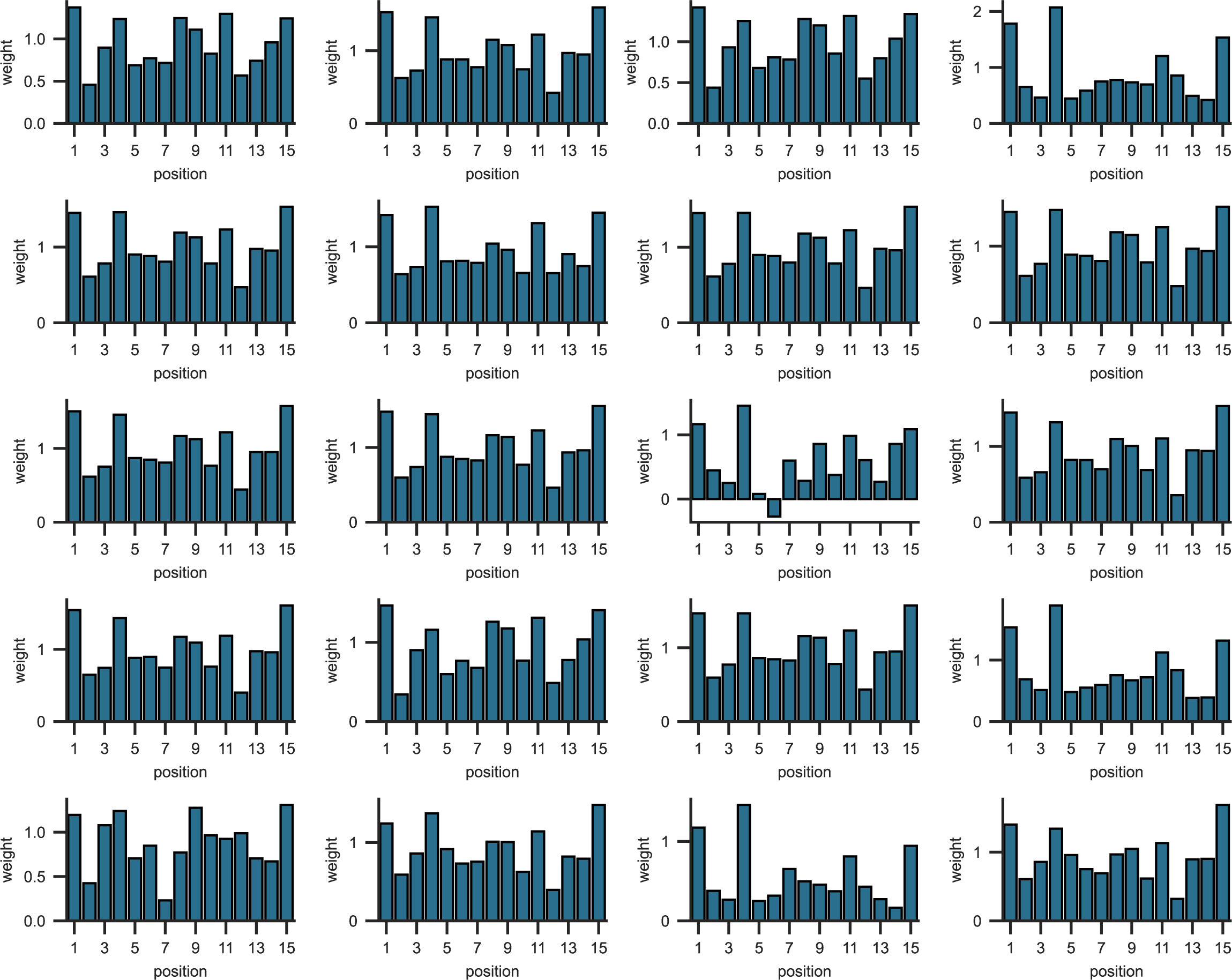
Kernel examples from “grouped” model training repetitions. The “grouped” has undergone 20 training runs with random initiations, and the resulting learned kernels are displayed.

**Supplementary Figure 6.**
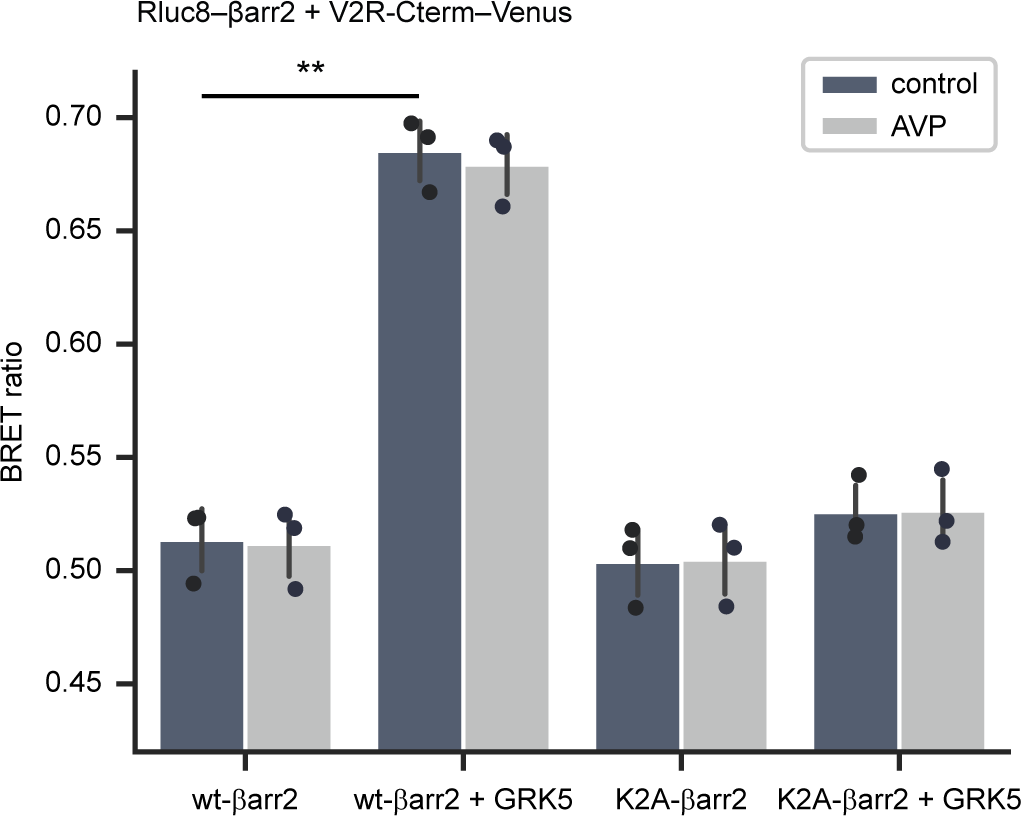
The core region of V2R is not necessary for βarr2 binding. BRET measurements between membrane-targeted V2R-Cterm–Venus andꞵwt-βarr2–Rluc8 or K2A-βarr2–Rluc8 in the absence or presence of GRK5 coexpression, a kinase with constitutive activity. The cells were either stimulated with vehicle or 100 nM AVP and raw BRET ratios are shown. Mean values ±S.E.M. are shown, and the data were statistically tested with one-way ANOVA, (n=3), **: p<0.0001.

**Supplementary Figure 7.**
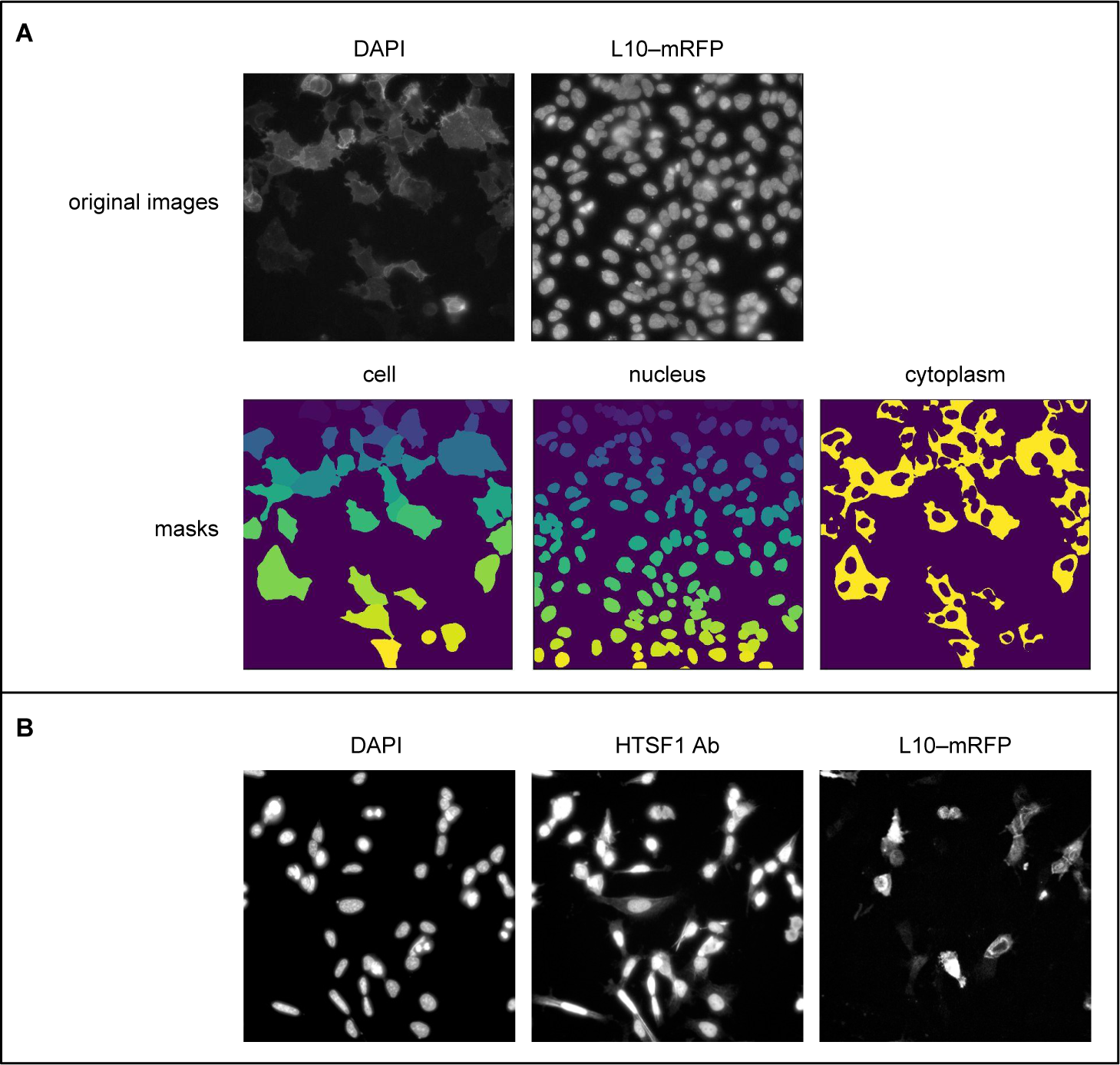
HTSF1 intracellular distribution. A. Quantification method of the intracellular location of the HTSF1. Images of DAPI-stained cells expressing L10-mRFP were taken with ImageXpress confocal microscope, the individual cell- and nuclear masks (bottom row) were detected with the Cellpose algorithm, and the cytoplasm masks were created by subtracting the nuclear mask from the cell mask. The masks were used for the determination of the HTSF1 fluorescence (mNeonGreen or HTSF1 staining) in the cytoplasm and nucleus. B. Representative confocal images of βarr1/2 KO HEK 293A cells expressing L10–mRFP. The cells were fixed, permeabilized, and stained with anti-HTSF1 antibody and DAPI. The images were gamma corrected with a value of 0.5 for better visualization of the low cytoplasmic HTSF1.

